# Functional reorganization of large-scale brain systems underlies emotion regulation maturation

**DOI:** 10.64898/2026.09.22.752555

**Authors:** Rui Ding, Zhi Zhang, Ze Zhang, Ruoting Qing, Bowen Hu, Qi Huang, Yi Luo, Yi Hu, Xiaolin Zhou, Shaozheng Qin

## Abstract

Emotion regulation via cognitive reappraisal is essential for socioemotional development and mental health, yet the organizational logic of its supporting brain systems across youth remains unclear. Using a large multisite cohort, we demonstrate that maturation of reappraisal capacity reflects reorganization of a distributed architecture rather than progressive strengthening of prefrontal control. Reappraisal robustly reduced negative affect, with non-linear gains concentrated between 6 and 10.5 years. Bayesian systems identification resolved this architecture into four functionally distinct systems—reappraisal-specific, common-appraisal, modifiable-emotion, and non-modifiable-emotion components—spanning prefrontal-control, salience, default-mode and dorsal-attention networks. Age-related variation in anterior temporal cortex, emerged as essential hub, statistically accounted for developmental gains in reappraisal success. Multivariate modelling together with intersubject analyses further identified a cross-site-generalizable whole-brain reappraisal signature and revealed an age-dependent shift in brain—behaviour coupling, with neurobehavioural convergence increasing with age. These findings establish distributed network reorganization as a core mechanism of emotion regulation development and provide mechanistic targets for stage-matched interventions.

## Introduction

Emotion regulation is essential to adaptive behavior, socioemotional development and mental health. Its aberration is implicated across various psychiatric disorders and is associated with both internalizing and externalizing symptoms in children and adolescents, making emotion dysregulation a major transdiagnostic dimension of developmental psychopathology ^1–4^. Cognitive reappraisal—the reinterpretation of an emotional event to alter its affective impact—is a central form of deliberate emotion regulation ^5,6^. Childhood and adolescence are particularly important periods for understanding this capacity, as rapid cognitive and socioemotional changes coincide with the emergence of many mental health problems ^7^. Yet when reappraisal capacity changes most reliably across youth, and how its neural substrates are organized across age, remain unclear. Resolving these questions is critical for understanding how adaptive emotion regulation is established and why vulnerability to dysregulation varies across development. A key question is whether age-related differences in reappraisal reflect the progressive strengthening of regulatory control or stage-sensitive reorganization of a distributed, multicomponent neural architecture.

Cognitive reappraisal is a multicomponent capacity whose constituent processes may follow different age-related trajectories. Successful reappraisal draws on attentional shifting, working memory, cognitive control, meaning reconstruction, affective processing and social understanding, each of which changes substantially but not synchronously across childhood and adolescence ^8–11^. The overlapping-waves theory provides a useful account of how such asynchronous changes can shape behavior: developmental change reflects shifts in the relative availability and efficiency of multiple coexisting processes rather than uniform improvement in a single capacity ^12^. Accordingly, as different component processes become more or less effective across age, their changing contributions may produce periods of rapid improvement, relative stability or other nonlinear variation in overall reappraisal performance. Consistent with this possibility, previous studies have reported heterogeneous age associations in reappraisal performance and habitual strategy use, rather than a single monotonic trajectory ^11,13^. We therefore hypothesize that reappraisal success would vary nonlinearly with age, revealing age-sensitive developmental windows in which change would be particularly pronounced.

This multicomponent account also has implications for the neural organization of cognitive reappraisal. Classical neurocognitive models emphasize lateral prefrontal and cingulate systems that maintain regulatory goals and exert top-down control over emotion-generative responses ^14,15^. Meta-analytic and systems-level studies, however, show that reappraisal engages a broader architecture spanning prefrontal, parietal, temporal, insular, midline and subcortical regions involved in cognitive control, appraisal, meaning reconstruction and affective processing ^16–18^. Developmental models of the emotional brain further propose that age-related differences in emotion regulation reflect changing organization across cortical and affective systems rather than prefrontal maturation alone ^19^, consistent with the broader principle of interactive specialization that functional development involves functional integration and specialization across distributed cortical systems ^20,21^. Importantly, distributed recruitment is not functionally homogeneous. Systems-identification work in adults has dissociated reappraisal-specific and common-appraisal activity from emotional responses that are either modifiable or relatively unchangeable emotion during regulation, revealing distinct functional components that span canonical network boundaries ^22^. If these components contribute differently to reappraisal and vary differently with age, age-related neural differences should extend beyond changes in prefrontal activation to the functional organization of the broader reappraisal architecture. We thus hypothesize that age-associated neural variation would involve multiple functionally distinct components and that these components would show differential associations with regulatory success.

A further question concerns how individual differences in reappraisal are organized across age. Developmental neuroimaging has traditionally emphasized age-related differences in mean behavior and regional activation but mean-level analyses cannot capture how individuals are organized relative to one another. Similar behavioral outcomes may be supported by different neural configurations, and the correspondence between neural and behavioral variation may itself differ across age. Recent work has shown systematic age-related changes in inter-individual similarity of socioaffective representations, including greater convergence among older youths ^23^. This population-level perspective therefore complements conventional trajectory analyses by asking whether behavioral and neural variability acquire a different relational structure across development. We thus asked whether inter-individual organization of reappraisal-related behavior and neural responses, and the correspondence between them, varied systematically with age.

To test the above hypotheses, we leveraged a large multisite sample of 1,316 participants aged 6–18 years to investigate brain systems involved in cognitive reappraisal across three complementary levels of organization. At the developmental level, we characterized nonlinear age-related variation in reappraisal success and regional task-evoked activity to identify when and where age differences were most pronounced. At the systems level, we combined Bayesian systems identification with cross-site multivariate modelling to determine how distributed reappraisal activity is functionally differentiated and integrated into a reproducible whole-brain representation. At the population level, we implemented intersubject similarity to test whether the organization of behavioural and neural individual differences, and their correspondence, varied systematically with age. By integrating developmental trajectories, functional architecture and population-level organization, we tested whether age-related variation in reappraisal is better characterized by coordinated changes across distributed brain systems than by uniform strengthening of regulatory control. This approach enables us to distinguish changes in response magnitude from changes in functional and inter-individual organization, linking when reappraisal varies with age, where those differences are expressed in the brain, and how neural variation relates to regulatory behavior across youth.

## Results

### Emotion regulation via cognitive reappraisal robustly reduces negative feelings in youth

We first examined the effectiveness of cognitive reappraisal in a cross-sectional sample of 1,316 youth aged 6-18 years recruited across five areas covering 8 sites (see *Supplementary Figure 1* for details). Participants viewed a total of 42 peer-like faces under three cognitive manipulations: (i) Reappraise: actively reappraising a negatively emotional event indicated by facial expression (fear/anger/sadness/disgust), (ii) Look: passively viewing a negative face, and (iii) Neutral: passively viewing a neutral face, followed by a self-report of their emotional response (**Fig. 1a**). The age distributions did not differ significantly between male and female participants or across sites (pairwise *ps* > 0.05; **Fig. 1b**).

**Fig. 1.**
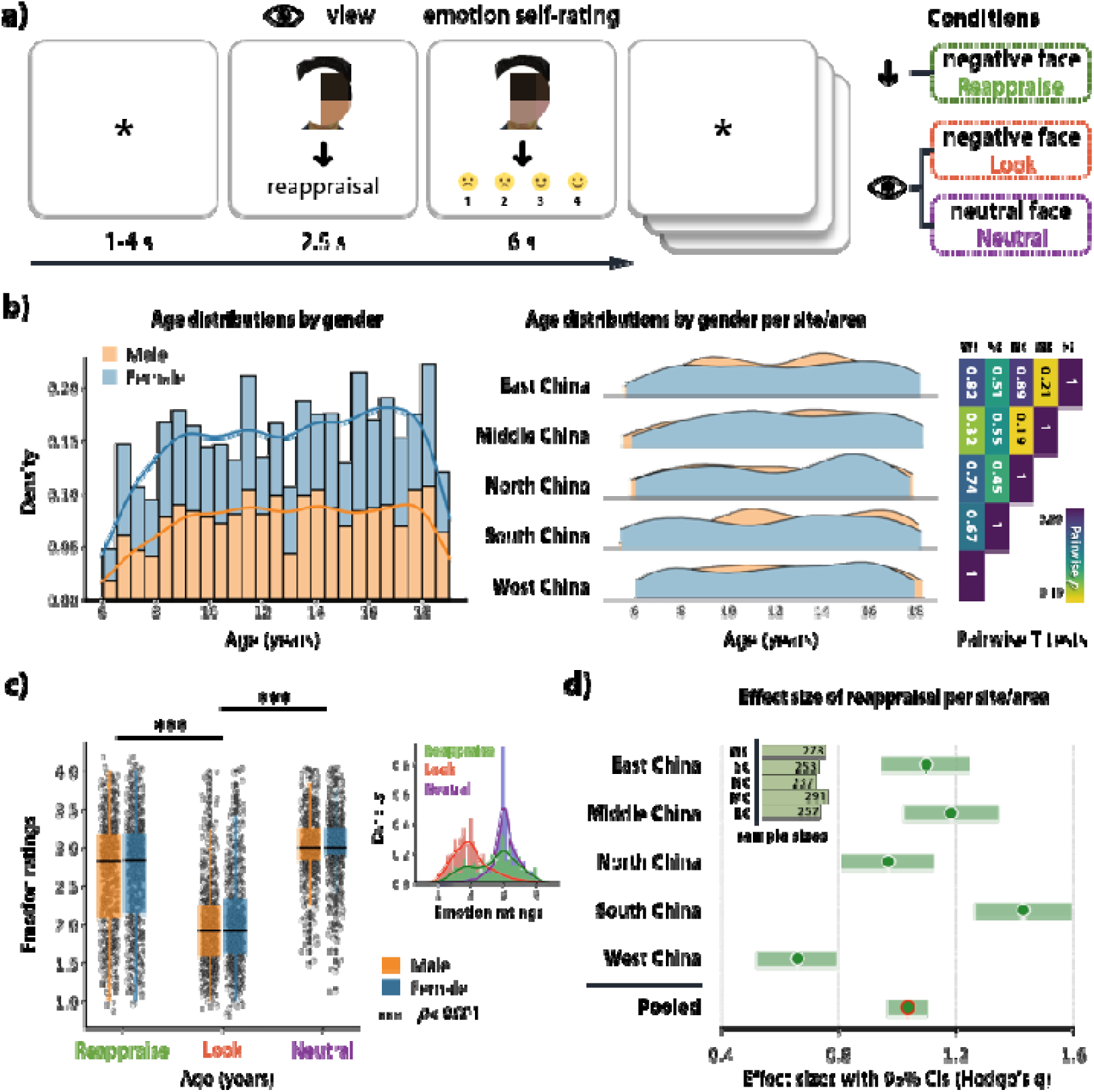
Experimental paradigm along with cross-sectional samples and behavioral effects. (**a**) The experimental paradigm of emotion regulation (cognitive reappraisal) used in the current cross-sectional samples. (**b**) The age distributions of samples show no significant difference between genders and across sites/areas (pairwise *ps* > 0.05). (**c**) The reappraisal condition manifests significant improvement of emotion feelings than passive viewing condition (Look), which is also confirmed in each site/area (**d**).

As shown by **Fig. 1c**, the task elicited robustly emotional and regulatory effects. Repeated measures ANOVA with cognitive manipulations as within-subject factors and sex as between-subject factor revealed significant effect across three conditions (F(2, 2628) = 1459.41, *p* < 0.001, partial η^2^ = 0.53). Yet, the sex effect on emotional self-rating did not reach significance (F(1, 1314) = 1.08, *p* = 0.30, partial η^2^ = 0.001). Passive viewing of negative facial expressions relative to neutral ones significantly altered emotional ratings, confirming robust responses to negative cues (post-hoc T = - 65.47, *p_holm_* < 0.001, Cohen’s d = - 1.81; **Fig. 1c**). Cognitive reappraisal significantly reduced negatively emotional ratings relative to passive viewing of the negative ones (post-hoc T = 32.73, *p_holm_* < 0.001, Cohen’s d = 0.90; **Fig. 1c**). This regulatory effect was consistently observed across all sites: site-specific effect estimates were positive, their confidence intervals excluded zero, and the pooled effect was approximately one standard deviation (Hedges’g = 1.03; **Fig. 1d**). Hence, cognitive reappraisal yields a robust and geographically replicable reduction in negative affect across the full age range, providing a basis for characterizing age-related variation in reappraisal success.

### Emotion regulation ability via reappraisal shows age-related nonlinear variation in youth

Building on above results of robust reappraisal effect in emotion regulation across youth, we next examined developmental changes in reappraisal success from 6 to 18 years old. Reappraisal success was defined as the difference in emotionally negative ratings between the cognitive reappraisal minus passive viewing negative event conditions (i.e. Reappraise - Look). Furthermore, we fitted generalized additive models (GAMs) to characterize potentially nonlinear age-related variation. The first derivatives of the posterior trajectory estimates were used to identify turning points across development and also the intervals during which the rate of age-related change differed from zero (**Fig. 2a**).

**Fig. 2.**
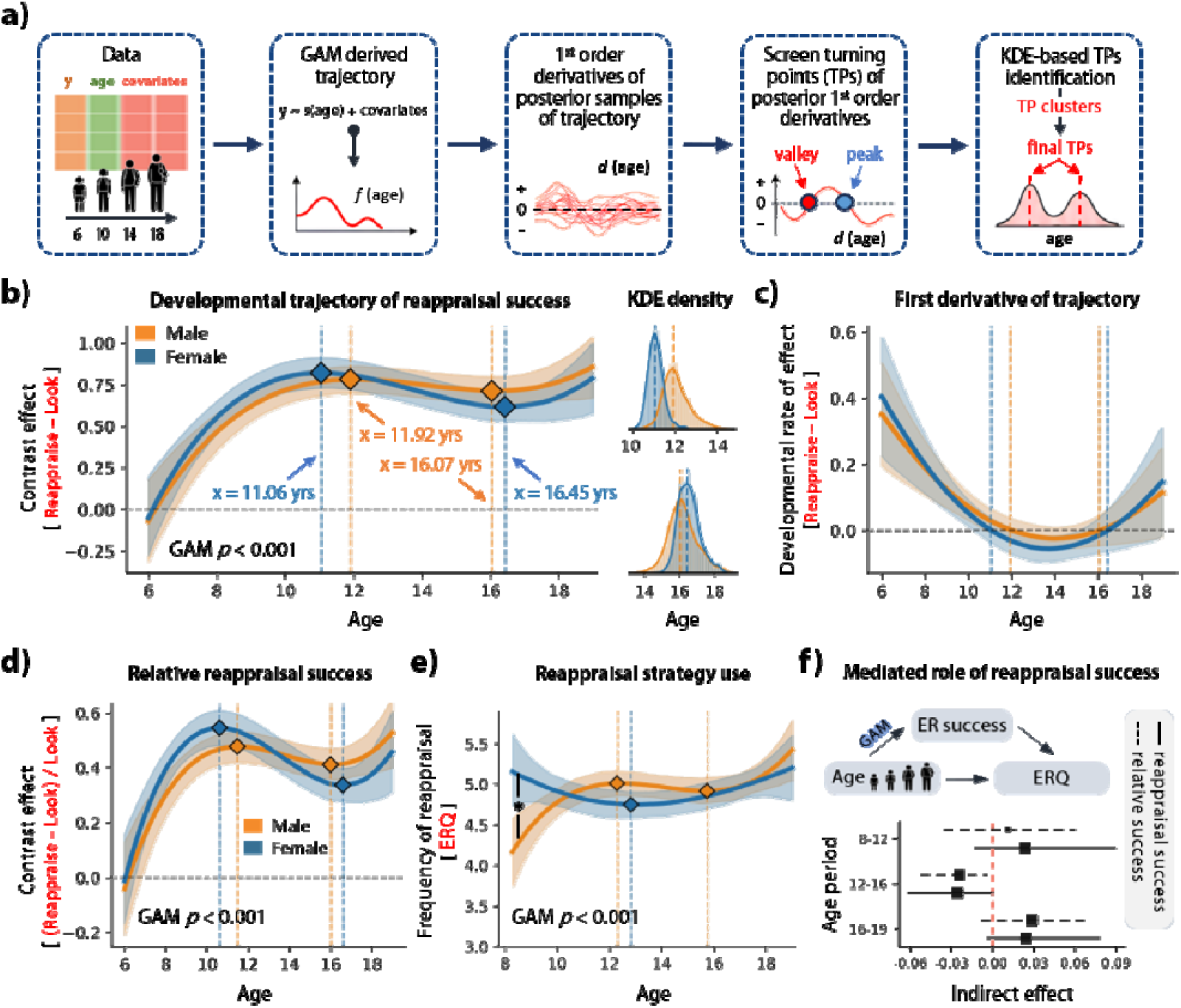
Age-related developmental changes in emotion regulation ability via reappraisal. (**a**) The procedure of trajectory fitting and identifying turning points. (**b**) The gender-specific developmental changes in reappraisal success is of significant association with age (*p_GAM_* < 0.001). And developmental rate of reappraisal ability across youth is significant away from zero during 6 – 10.5 years period (**c**). The relative reappraisal success was also computed, showing significant age-related trajectory (*p_GAM_* < 0.001) which has similar shape with reappraisal success (**d**). The reappraisal strategy use measured by ERQ is significantly associated with age (**e**), and (**f**) reappraisal success significantly mediated the association between age and reappraisal strategy use during 12 - 16 years period (reappraisal success: indirect effect = -0.025, *p* = 0.018; relative reappraisal success: indirect effect = -0.024, *p* = 0.002).

GAM analysis revealed a significantly non-linear age-related changes in reappraisal success from childhood through adolescence into adulthood (pseudo R^2^ = 0.045, effective degree of freedom (EDoF) = 7.28, *p_GAM_* < 0.001; **Fig. 2b**). The fitted trajectories increased steeply from 6 to 10.5 years old during middle childhood and reached an initial turning points at 11.06 years in females and 11.92 years in males. The trajectories subsequently showed a modest decline, with later turning points estimated at 16.45 years in females and 16.07 years in males, followed by an increase toward the upper end of the sampled age range (**Fig. 2b**). Further derivative analysis provided statistically reliable evidence for age-related changes only during the earlier portion of the trajectory: indicated by 95% confidence intervals of the first derivatives of developmental trajectories, the developmental rate was significantly positive from approximately 6 to 10.5 years (males: 10.77 years, females: 10.24 years), whereas the later negative and positive portions of the fitted trajectories did not differ reliably from zero (**Fig. 2c**). Thus, the strongest inferential evidence was for developmental gains in reappraisal success during middle childhood, whereas the later fluctuations represented features of the fitted cross-sectional trajectory that were not independently supported by significant derivative intervals.

To determine whether this developmental pattern is driven by baseline reactivity to negative emotion stimuli, we additionally calculated the relative reappraisal success by normalizing the reappraisal effect to emotional responses during passive viewing of negative stimuli. GAM analysis using relative reappraisal success as dependent variable also revealed significant age-related variations (pseudo R^2^ = 0.037, EDoF =7.28, *p_GAM_* < 0.001) and showed a trajectory similar to that of the absolute reappraisal-success measure (**Fig. 2d**). Habitual tendency of reappraisal use, assessed using the Emotion Regulation Questionnaire (ERQ), was also significantly associated with age (pseudo R^2^ = 0.02, EDoF = 7.74, *p_GAM_* < 0.001), although its fitted trajectory differed from the task-based measures of reappraisal success (**Fig. 2e**).

To further test whether actual reappraisal ability accounted for age-related variation in habitual reappraisal use over development, we conducted mediation analyses using task-based reappraisal success as mediator linking age and reappraisal strategy use (upper panel of **Fig. 2f**). The path linking age and reappraisal success was estimated using GAM. These analyses revealed significant indirect effects specifically during 12-16 years for both absolute reappraisal success (indirect effect = -0.025, 95%CI = [-0.06, -6.46×10^-6^], *p* = 0.018) and relative reappraisal success (indirect effect = -0.024, 95%CI = [-0.05, -0.004], *p* = 0.002). Such indirect effects, however, were not significant in younger or older age periods (lower panel of **Fig. 2f**). These results establish a nonlinear age-related behavioral phenotype over development, with the strongest gain in reappraisal success during middle childhood and an age-specific statistical association between reappraisal task performance and habitual strategy use for emotion regulation during adolescence.

### Emotion regulation via cognitive reappraisal recruits distributed brain networks and nonlinear developmental changes across youth

We next identify brain systems and networks involved in emotion regulation via cognitive reappraisal collapsing across age range. Reappraisal-related activation was defined by the contrast between reappraising negative stimuli and passively viewing negative ones (i.e. Reappraise - Look), whereas emotional reactivity was defined by the contrast between passively viewing negative stimuli and passively viewing neutral ones. Both contrasts elicited robust activation in a network of widely distributed cortical and subcortical regions (FDR *q* < 0.005; **Fig. 3a**), encompassing lateral prefrontal, temporoparietal junction, medial prefrontal, precuneus and cerebellar regions (*Supplementary Table 1*). For reference, we also calculated the brain activations during emotional reactivity. Emotional reactivity recruited a network of partly overlapping yet dissociable regions spanning lateral prefrontal and insular cortex, medial frontal and cingulate regions, superior temporal cortex, thalamus, amygdala and cerebellum (**Fig. 3a** and *Supplementary Table 2*).

**Fig. 3.** 

The spatial relationship between the two contrasts differed across cortical and subcortical systems. Reappraisal and reactivity maps were nearly orthogonal in the cortex (r = - 0.03) but showed a stronger inverse spatial relationship in subcortical regions (r = - 0.32). The whole brain activation was parcellated based on Schaefer parcellation atlas ^24^, and subsequent region-level estimates showed a resulting pattern similar to the voxel-wise estimation (middle panel of **Fig. 3a**). Both youth activation maps demonstrated modest spatial correspondence with independent meta-analytic (reappraisal: r = 0.56, reactivity: r = 0.48) and brain component maps (reappraisal: r = 0.28/0.42, reactivity: r = 0.29 for Neurosynth, r = 0.10 for Morawetz) of emotion regulation and emotional reactivity derived from Neurosynth dataset and Morawetz et al. (2020) ^18^ (**Fig. 3b**). This indicates a plausible emergence in youth of the neural activity patterns of emotion generation and regulation collapsing across all ages. At the network level, significant reappraisal-related activation was distributed across canonical cortical systems ^25^, with relatively prominent contributions from the default-mode and dorsal-attention networks compared with emotional reactivity (**Fig. 3c**). Altogether, regulating emotion elicited by negative faces recruits similar brain regions to conventional emotion regulation, and was not localized to a single control network but was expressed across multiple large-scale neural systems.

Importantly, we examined the association of reappraisal-evoked activation with age using both GLM and GAM. Consistent with the nonlinear age-related variation observed for behavioral reappraisal success, GAM analysis for brain activation using the same parameters also identified significantly nonlinear age effect (FDR *q* < 0.05), with most prominent effects covering the dorsal lateral prefrontal cortex, insula, temporal, default-mode regions, temporoparietal junction area, middle cingulate cortex and precuneus (regions outlined in red in **Fig. 3d**, see *Supplementary Figure 2* for more detailed trajectories and corresponding brain regions). GLM analysis, however, did not identify any regions surviving multiple comparisons correction. To complement above regional analyses, we characterized age-binned changes in the spatial organization of reappraisal-related neural responses based on brain mappings of emotion regulation established in previous neuroimaging meta-analysis ^18^ and in Neurosynth dataset ^26^ serving as a reference in adults brains. One hand, spatial similarity with both independent emotion-regulation maps fluctuate across age bins rather than changing monotonically, in which the similarity increases from approximately 6 to 11 years and then fluctuates at a relatively stable levels until 19 years (left panel of **Fig. 3e**). On the other hand, indexed by proportion of the brain map’s total significant effects that fall within a given network, we decomposed the network contribution to reappraisal-elicited activation. Further network decomposition likewise revealed changes in the relative composition of the reappraisal neural response, including a descriptive increase in the relative contribution of the default-mode network and decrease in visual and dorsal-attention network toward older age bins (right panel of **Fig. 3e**). Together, these analyses indicate that emotion regulation via cognitive reappraisal recruits a widely distributed neural architecture within which activation magnitude and spatial organization vary from 6 to 18 years old in youth.

### Bayesian systems identification of brain systems involved in emotion regulation and emotion reactivity in youth and their developmental effects

As a complex construct of mental processes, emotion regulation via cognitive reappraisal in socioaffective circumstance recruits multiple brain networks which is functionally heterogeneous activation profiles ^22^. We therefore partitioned reappraisal-related brain activity maps using the joint Bayes factors of effects across emotion regulation (Reappraisal condition versus passive viewing negative condition) and emotion reactivity (passive viewing negative condition versus passive viewing neutral condition) contrasts (**Fig. 4a**). Based on Bayes factors, this axiomatic systems-identification method successfully distinguished four components: a reappraisal-specific component activated by emotion regulation but not emotion reactivity; a common-appraisal component activated by emotion reactivity and further recruited during emotion regulation; a non-modifiable emotion component activated by emotion reactivity but not altered by emotion regulation; and a modifiable emotion component activated by emotion reactivity and reduced during emotion regulation **(Fig. 4a**).

**Fig. 4.**
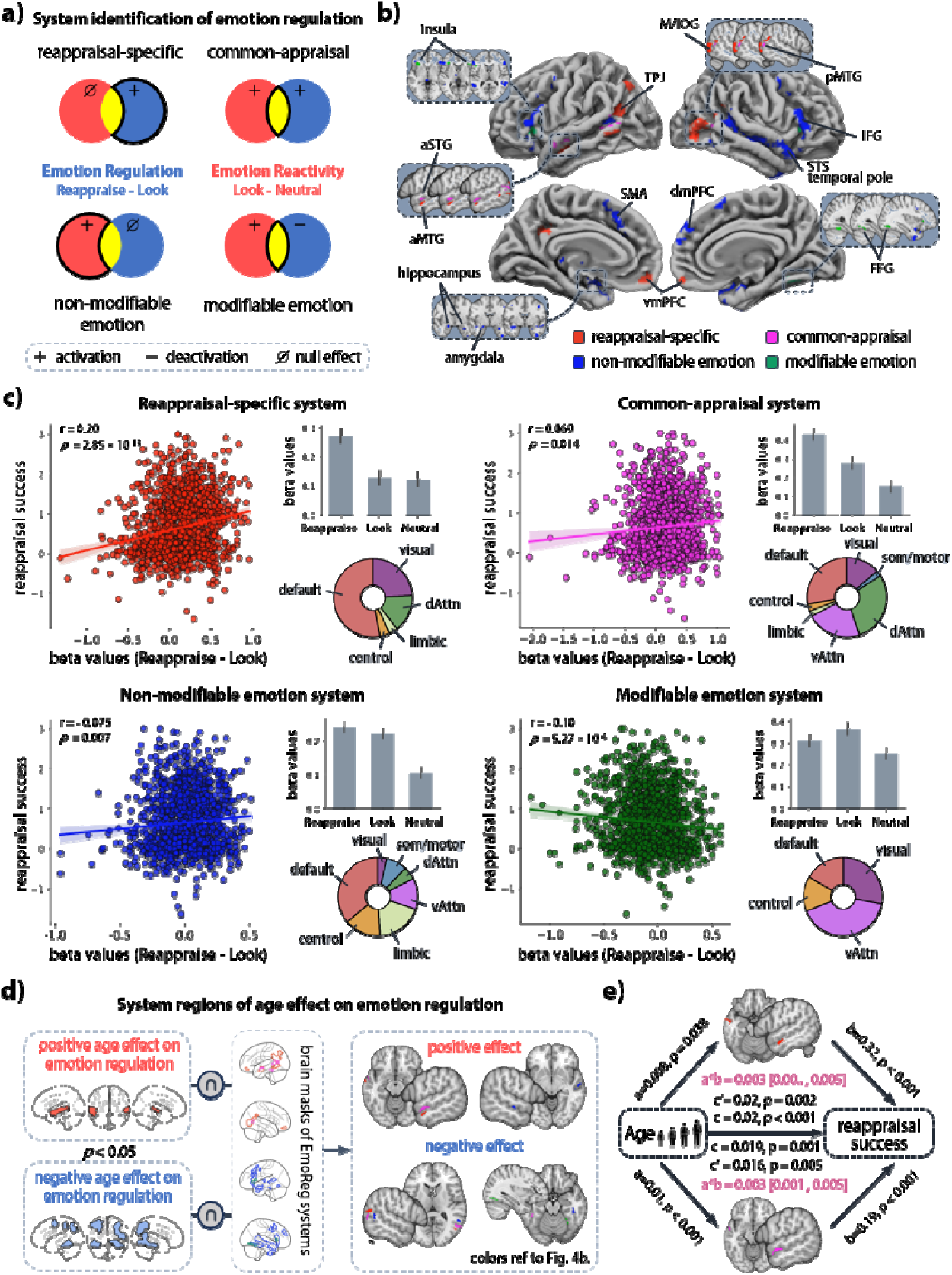
Reappraisal-specific and common-appraisal brain systems along with non-modifiable and modifiable emotion components in youth and corresponding developmental effects. (a) The logic for identifying brain systems of emotion regulation is represented in Venn diagrams, in which the circles represent voxels of emotion regulation (reappraisal) and reactivity. The proportion of the Venn diagrams matching each brain system is outlined in black. (b) The brain regions corresponding to each brain systems are projected to brain surface and detailed in slices using different colors. (c) The associations between brain activation within brain systems and reappraisal success are depicted in scatter plots, along which the network contributions to each brain system and the activation patterns matching the logic of systems identification are visualized. (d) We localized system regions of age effect, (e) and found the significant mediated roles of aSTG (common appraisal) and aMTG (reappraisal only) in age-dependent increase of reappraisal success.

All four components were spatially distributed and extended across multiple canonical networks (**Fig. 4b** and pie charts of **Fig. 4c**). The reappraisal-specific component showed a strong contribution from the default-mode network, together with weaker contribution from visual and dorsal-attention networks, and weakest contribution from control and limbic networks. The common-appraisal component was evenly distributed across default-mode, visual, dorsal- and ventral-attention networks, and also minorly distributed across control, somatomotor and limbic networks. The non-modifiable-emotion component predominantly covered default-mode, control, limbic and ventral-attention networks, whereas the modifiable-emotion component was concentrated primarily in ventral-attention, visual, control and default-mode networks (pie charts in **Fig. 4c**). In sum, functional component membership and canonical network membership provides complementary descriptions of the neural organization of cognitive reappraisal: each component of functional profiles of emotion regulation is itself implemented across multiple large-scale networks, which keeps consistent with findings identified in adults ^22^.

The four components in youth also differ in their associations (i.e. Pearson correlations) with individual differences in reappraisal success. Greater activation of reappraisal-specific system was positively associated with reappraisal success (r = 0.20, *p* = 2.85×10^−13^), representing the strongest component-level association (**Fig. 4c**). Activation in common-appraisal and non-modifiable-emotion component respectively showed weaker positive associations with reappraisal success (former: r = 0.069, *p* = 0.014; latter: r = 0.075, *p* = 0.007; **Fig. 4c**). In contrast, the activation in modifiable-emotion component was positively associated with reappraisal success (r = − 0.10, *p* = 5.27×10^−4^; **Fig. 4c**). These distinct brain-behavior associations, together with different condition-response profiles (bar charts of **Fig. 4c**), further support that these partitioned components represent functionally dissociable neural subsystems, each playing a specialized and distinct role in emotion regulation rather than operating as a single, homogeneous regulatory network.

We next situated the age-activation associations identified in the preceding whole-brain analysis within this functional architecture. Positive and negative age-associated activation regions (FDR *p* < 0.05) were intersected with the four component masks, yielding candidate age-associated regions within the functional architecture (**Fig. 4d**). These conjunctions were used to define regions for subsequent mediation analyses using brain activation mediator linking age variation and individual differences in reappraisal ability. The mediation analysis demonstrated that the anterior middle temporal gyrus (aMTG) within the reappraisal-only component significantly mediated the association between age and reappraisal success (indirect effect = 0.003, 95% CI = [0.001, 0.005]). In the anterior superior temporal gyrus (aSTG) within the common-appraisal component, we also identified the mediated role of neural activity in association between age and reappraisal success (indirect effect = 0.003, 95% CI = [2.88×10^-5^, 0.005]). The direct age-reappraisal success association remained significant in both models (**Fig. 4e**). Thus, youth reappraisal in reappraisal-only and common-appraisal systems statistically accounted for part of the cross-sectional association between age and reappraisal success, suggesting that age-related improvements in cognitive reappraisal ability are driven by functional maturation of these functionally dissociable brain subsystems.

### Multivariate activity during reappraisal forms a cross-site generalizable neural signature across youth

The preceding analyses characterized emotion regulation via cognitive reappraisal recruited a set of widely distributed brain systems and networks at task-invoked regional activation, network and dissociable component levels. To further test whether such distributed activation could be integrated into a generalizable neural representation, we trained multivariate decoding models to distinguish cognitive reappraisal from passive viewing of negative faces using leave-one-site-out validation procedure. Model optimization was performed within each training set (**Fig. 5a**). All three classifiers distinguished reappraisal from passive viewing significantly better than chance in held-out sites. Logistic L2 regression achieved an area under the receiver operating characteristic curve (AUC) of 0.69, linear support vector machine (SVM) achieved an AUC of 0.70, and radial-basis-function SVM achieved an AUC of 0.69 (left and middle panels of **Fig. 5b**). All model performances reached statistical significance in permutation tests (permuted *ps* < 0.001; right panel of **Fig. 5b**). A similar performance of linear and nonlinear models indicated that the generalizable condition information was captured adequately by a predominantly linear multivariate boundary. We therefore characterized the distributed neural signature using the linear model in the subsequent analyses. To identify significant robust brain regions (i.e. features) contributing to classification, the whole samples were bootstrapped 10000 times with replacement, and refitted the multivariate model. Based on derived bootstrapped weight distributions, the region-wise two-sided p values were estimated. The results revealed the significant regions contributing to Reappraisal prediction, mainly covering dorsal lateral prefrontal cortex, temporal default-mode regions, temporoparietal junction and precuneus (FDR *q* < 0.05; **Fig. 5c**). The ventral lateral prefrontal cortex, insula and medial prefrontal cortex preferentially contribute to a robust prediction of passive viewing negative condition (FDR *q* < 0.05; **Fig. 5c**). Expression of this integrated signature showed statistically significant age-related changes across youth. GAM analysis explained 1.0% of the variance in signature response (R^2^ = 0.01, *p* = 5.31×10^−8^), whereas the corresponding linear model explained 0.8% (R^2^ = 0.008, *p* = 0.015; Fig. 5d). Thus, age was reliably associated with expression of the multivariate reappraisal pattern, although the amount of variance explained was small.

**Fig. 5.**
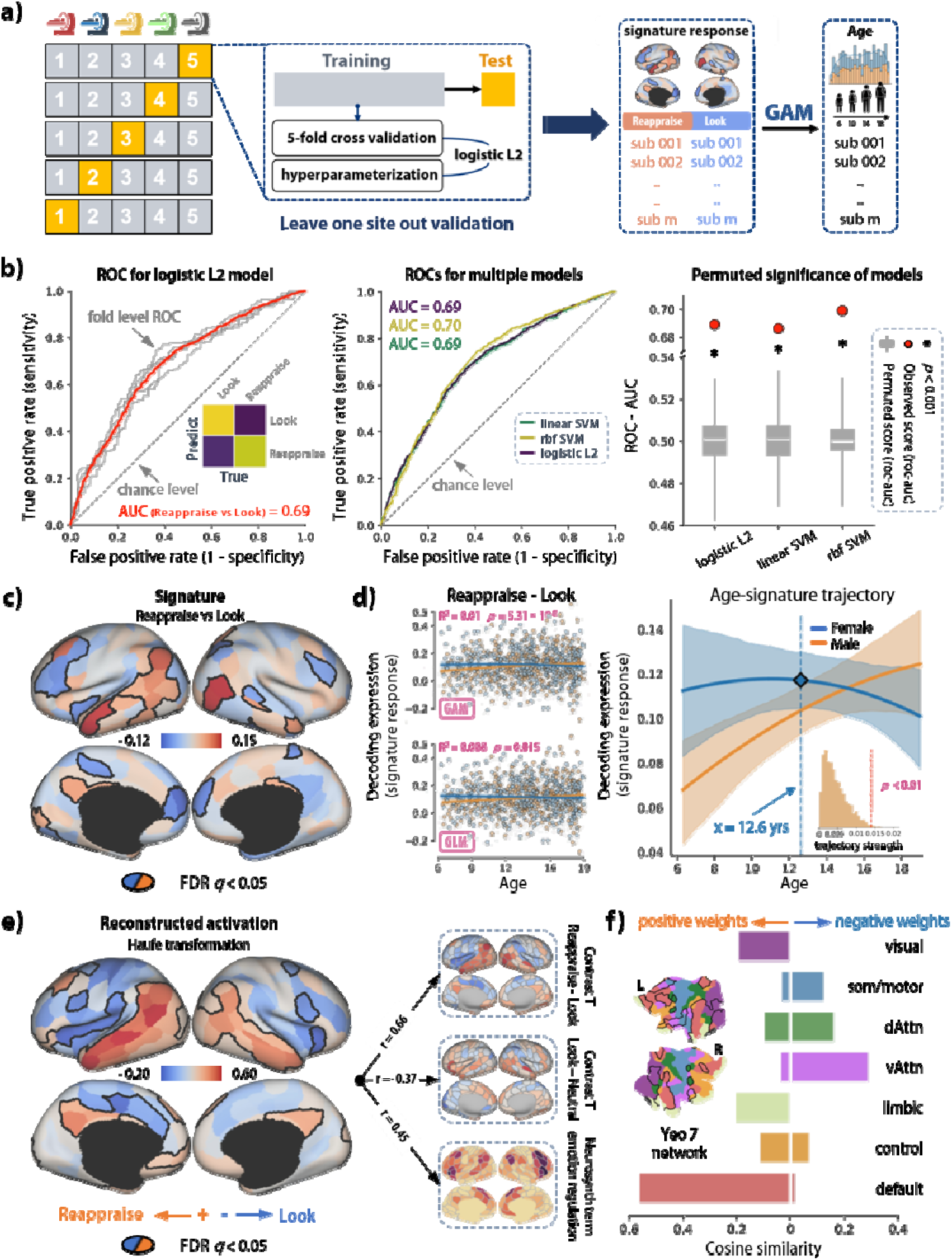
The multivariate neural signature of emotion regulation via cognitive reappraisal and its maturation across youth. (**a**) The procedure for multivariate model training and testing is followed by signature computation and GAM/GLM based association between neural signature of emotion regulation and age. (**b**) We used both linear (logistic L2 & linear SVM) and nonlinear models (rbf SVM) to learn the classification plane between reappraisal and passive view negative condition (i.e. Reappraise vs Look). And three models all reach statistical significance relative to chance level (permuted *ps* < 0.001). (**c**) The neural signature of reappraisal is projected onto brain surface (i.e. decoder weights map). The regions significantly contributing to the classification (Reappraise vs Look) are outlined in black (FDR *q* < 0.05), which were computed using bootstrapped *p* values of regions. (**d**) The association between neural signature response and age is simultaneously fitted using GAM and GLM and found significant association especially in male youth (GAM: R^2^ = 0.01, *p* = 5.31 × 10^-8^; GLM: R^2^ = 0.008, *p* = 0.015). (**e**) The reconstructed activation is computed through Haufe transformation, in which significant regions derived from bootstrapped decoder weights are outlined in black (FDR *q* < 0.05). The reconstructed activation of neural signature is significantly associated with univariate activation maps and emotion regulation mapping derived from Neurosynth dataset, and brain network contribution to neural signature is visually depicted in (**f**).

To improve interpretation of the discriminative pattern, we also applied a Haufe transformation to reconstruct activation associated with the signature ^27^.The reconstructed pattern showed high similarity with decoder weights map, and is strongly correlated with the univariate reappraisal contrast (r = 0.66) and with an independent Neurosynth-derived emotion-regulation map (r = 0.45; **Fig. 5e**), whereas showing an inverse spatial relationship with the emotional-reactivity contrast (r = − 0.37). Indexed by cosine similarity, neural signature of reappraisal showed distributed network contributions, with the default-mode, control, limbic and visual networks making the positive contribution and somatomotor and attention networks contributing prominently in the opposite direction (**Fig. 5f**). Together, these results indicate that youth aged 6-18 has already formed a cross-site generalizable neural signature that integrates distributed brain recruitments of reappraisal in socioaffective circumstance, although the classification performance did not reach as high level as the neural signatures derived from adults ^28^. Crucially, such a whole-brain signature exhibits a significant age-related increase in expression, indicating that the integrated, brain-wide neural network underlying cognitive reappraisal continues to mature, consolidate, and become more robustly engaged from childhood through adolescence.

### Developmental reorganization of brain systems reflecting inter-subject differences in emotion regulation ability

To address the central question of how brain networks involved in cognitive reappraisal reorganize and refine over development to support nuanced emotion regulation ability, we implemented an innovative approach through modeling the pattern of similarities and differences among individuals for single trial-invoked neural activity during cognitive reappraisal. Specifically, single-trial responses were aggregated by emotion category, and pairwise Mahalanobis distances were used to construct participant-by-participant behavioral and regional neural dissimilarity matrices (**Fig. 6a**). We compared these observed matrices with three models representing distinct developmental structures (see Developmental model in **Fig. 6a**): nearest neighbor, in which individuals closer in age are more similar; convergence, in which similarity increases toward older ages; and divergence, in which inter-individual differences increase toward older ages.

**Fig. 6.**
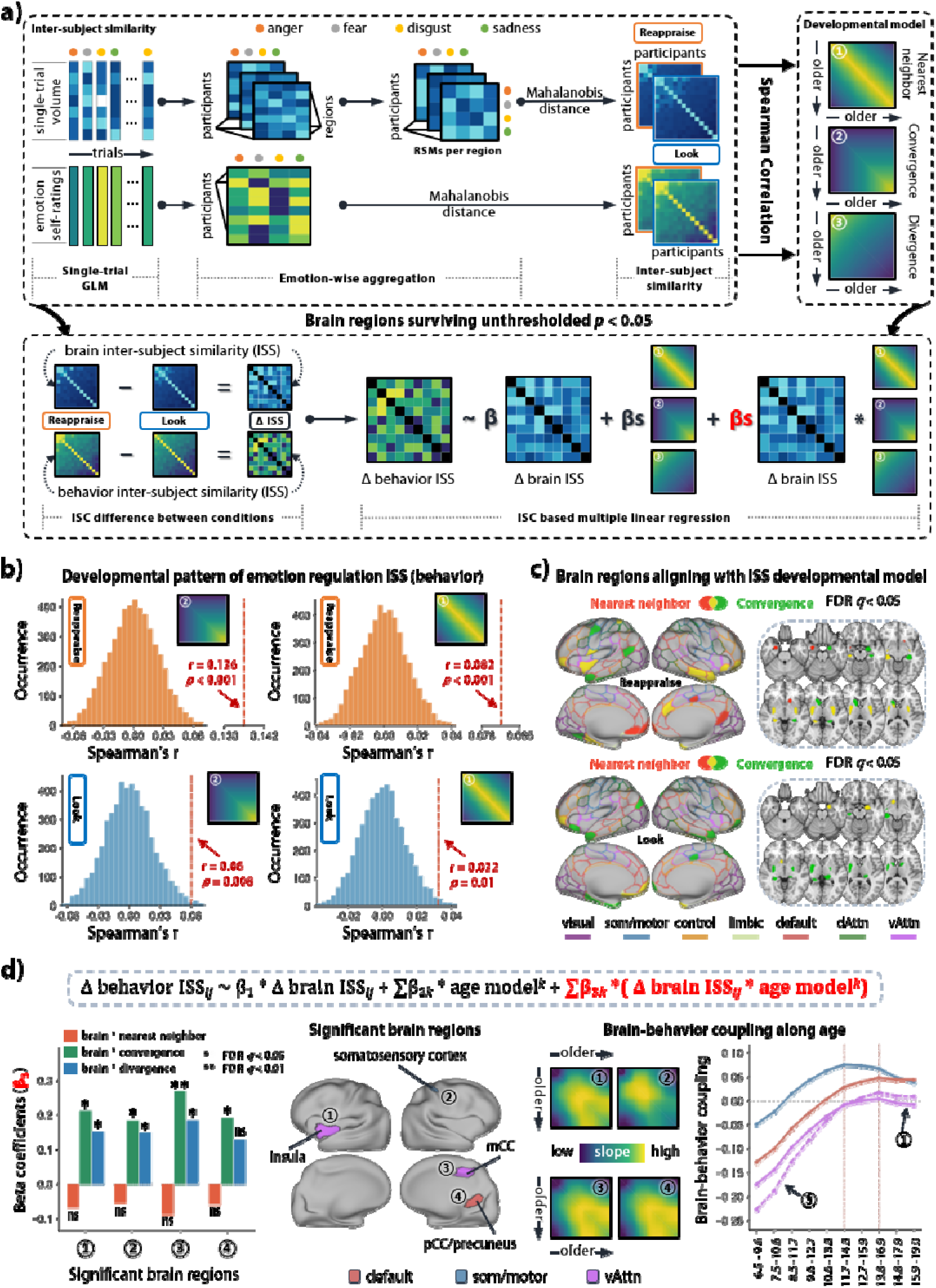
Inter-subject differences in brain systems underlying emotion regulation via cognitive reappraisal and their developmental reorganization. (a) A schematic chart depicts the procedure for model-based inter-subject similarity analysis to identify developmental pattern of inter-subject difference of reappraisal ability and corresponding brain basis. (b) The developmental pattern of inter-subject similarity in reappraisal response is significantly matching convergence model (CON) and nearest neighbor model (NN), the former of which has higher proximity with observations than the latter (CON: r = 0.136, *p* < 0.001; NN: r = 0.082, *p* < 0.001). (c) Brain regions where reappraisal-evoked activation significantly align with developmental models of inter-subject similarity are mapped onto brain surface and slices (FDR *q* < 0.05), with canonical brain networks outlined in corresponding colors. (d) The multiple linear model was conducted to identify brain regions underlying age-behavior association of inter-subject difference of reappraisal response. Four regions were successfully identified, in which the inter-subject similarity of reappraisal-evoked responses significantly moderated the age-behavior associations under developmental models of convergence and divergence. The brain-behavior coupling along age indexed by conditional slope is visualized in heatmaps, demonstrating that brain-behavior coupling is increasing until middle adolescence and then decreasing. This pattern is also identified in sliding window analysis (see rightmost panel of subplot d).

At the behavioral level, Mantel test with Spearman correlation revealed that inter-individual differences of reappraisal success were significantly associated with both convergence and nearest-neighbor models (**Fig. 6b**). The convergence model showed the stronger correspondence with the observed reappraisal matrix (r = 0.136, permuted *p* < 0.001), whereas the nearest-neighbor model also showed a significant but weaker association (r = 0.082, permuted *p* < 0.001). Passive viewing of negative faces showed substantially weaker developmental structure of nearest-neighbor model (r = 0.060, permuted *p* = 0.008) and convergence model (r = 0.032, permuted *p* = 0.010; **Fig. 6b**). In a word, the organization of individual differences was more pronounced during active reappraisal than during passive negative viewing and was predominantly characterized by greater similarity toward older ages, while retaining an additional contribution from developmental proximity.

Subsequently, we identified brain regions where neural inter-individual similarity aligned with the developmental models. Such alignment is indexed by Spearman correlation between inter-individual matrix of brain activities and age/developmental models, and the brain representations of these alignments are delineated in *Supplementary Figure 3*. Reappraisal-evoked neural responses showed significant nearest-neighbor and convergence structure across distributed regions spanning default-mode, dorsal-attention, ventral-attention, control, somatomotor, visual and limbic networks, especially ventral lateral prefrontal cortex, insula, temporal poles and hippocampus, caudate, putamen in subcortical regions (FDR *q* < 0.05; **Fig. 6c**; *Supplementary Figure 4* and *Supplementary Table 3*). Passive negative viewing also showed significant model-aligned regions, although the spatial pattern of cortex differed slightly from that observed during reappraisal (**Fig. 6c**). These results indicate that developmental organization of inter-individual differences is expressed not only in reappraisal behavior but also in regionally distributed neural response patterns.

Finally, we investigated whether the correspondence between neural and behavioral inter-individual differences itself varied across developmental position. For each candidate region derived from above developmental model-based analysis (*p* < 0.05), we modeled the difference of behavioral inter-subject similarity between reappraisal and passive viewing negative condition as a function of the corresponding neural-similarity difference, developmental-model terms and their interactions using multiple linear modeling (ISS-MLR; **Fig. 6d**). Four regions showed significant interactions between neural similarity and developmental structure: insula, somatosensory cortex, middle cingulate cortex and posterior cingulate cortex/precuneus (surface mapping in **Fig. 6d**). Neural-similarity interactions with the convergence model were significant in all four regions, interactions with the divergence model were significant in three regions, and nearest-neighbor interactions were not significant (bar charts in **Fig. 6d**). Furthermore, conditional-slope analyses under ISS-MLR revealed a common age-organized pattern across these regions. The association between neural and behavioral similarity was negative at younger ages, increased across childhood and early adolescence, crossed zero and became positive, reached its strongest positive values around middle adolescence, and then modestly attenuated toward late adolescence (heatmaps in **Fig. 6d**). Sliding-window analyses reproduced this overall pattern. Therefore, the predominantly convergent organization of behavioral and neural individual differences was accompanied by systematic age-related variation in their correspondence, showing that the behavioral relevance of neural similarity was not constant across the sampled age range. Altogether, these analyses demonstrated that age-related variation in reappraisal is expressed not only in average behavior, activation and multivariate pattern expression, but also in the developmental organization of inter-individual differences, highlighting that the maturation of emotion regulation involves a dynamic, qualitative reorganization of brain-behavior relationships, as individuals systematically converge toward a shared, mature neurobehavioral profile from childhood through adolescence.

## Discussion

By leveraging a multisite cross-sectional neuroimaging cohort of 1,316 participants aged 6-18 years, together with a cognitive reappraisal paradigm and complementary behavioral assessments, we characterized developmental reorganization of distributed brain systems underlying emotion regulation across youth. Behaviorally, cognitive reappraisal robustly reduced negative affect across all sites, while reappraisal ability indexed by reappraisal success showed nonlinear age-related developmental changes, with most prominent gains concentrated between approximately 6 and 10.5 years. At the regional level, reappraisal recruited a widely distributed set of cortical, subcortical and cerebellar regions, and age-related variation extended beyond canonical prefrontal control areas to temporal, insular, lateral temporal-parietal, and middle line regions. Bayesian systems identification further decomposed this distributed neural architecture into reappraisal-specific, common-appraisal, modifiable-emotion and non-modifiable-emotion components of distinct associations with regulatory success, which spatially overlapped with regions showing age-related variation and mediated the association between age and reappraisal success. At the whole-brain level, multivariate modelling identified a generalizable neural signature of reappraisal that generalized across held-out sites, and notably chronological age accounted for a small proportion of individual variation in its expression. Finally, intersubject-similarity analyses revealed that behavioral and neural response patterns during reappraisal became increasingly similar across individuals with increasing age, together with an age-dependent shift in neural-behavioral correspondence from negative towards positive associations across childhood and into mid-adolescence. Collectively, these findings suggest that cognitive reappraisal is supported by a distributed and functionally differentiated neural architecture that is already established across the sampled age range, whereas age-related differences are characterized by nonlinear variation in its regulatory effectiveness, regional and whole-brain functional organization and the population-level reorganization of brain-behavior relationship.

Behaviorally, reappraisal success showed the nonlinear age-related pattern. Participants throughout the sampled age range could successfully implement the instructed strategy, but reliable gains were concentrated between approximately 6 and 10.5 years rather than increasing uniformly across youth. This pattern is consistent with the substantial heterogeneity reported across developmental studies of reappraisal and argues against treating regulatory capacity as a unitary ability that changes at a constant rate ^11,13^. One interpretation for this developmental pattern follows the overlapping-waves theory framework: reappraisal depends on multiple capacities that do not vary synchronously with age, and changes in their relative contribution may therefore produce periods of accelerated improvement interspersed with relative stability ^12^. This interpretation remains indirect due to unavailability of measuring the constituent cognitive processes. Nevertheless, recent longitudinal evidence likewise indicates that explicit and implicit forms of emotion regulation can exhibit distinct regional age trajectories rather than a common maturational profile ^29^. The age-dependent correspondence between task performance and habitual reappraisal further underscores this distinction. As separable dimensions, the correspondence between regulatory capacity and regulatory tendency may itself vary with age ^30^, which is mirrored in our finding that task-based reappraisal success partially accounted for association between age and habitual reappraisal use only during mid-adolescence but not during other periods.

The neural findings provide a systems-level context for this behavioral nonlinearity. Reappraisal recruited a broad architecture encompassing prefrontal, cingulate, parietal, temporal, insular, subcortical and cerebellar regions, while age-related variation extended across dorsolateral prefrontal, insular, temporal, temporoparietal, middle cingulate and precuneus regions. Notably, age effects were distributed over multiple networks, arguing against a simple account of progressively stronger prefrontal recruitment across youth. In contrast, this distributed developmental pattern is consistent with our hypothesis that age-related differences in reappraisal involve coordinated variation across multiple functionally distinct neural systems ^19^. This interpretation also accords with contemporary accounts of reappraisal as a multicomponent process ^22,28^. Reappraisal does not simply suppress a generated emotional response, and it also alters how an event is construed or evaluated and thereby changes its affective meaning ^22,31,32^. From this perspective, age-related variation spanning multiple networks may therefore reflect development not only in regulatory control, including how emotional information is appraised and reframed, but also in how emotional information is represented ^33^. Although the present findings do not assign specific computations to individual regions, they indicate that age-related variation spans both regulatory operations and the representations on which those operations act.

The systems-identification results further show that distributed reappraisal activity is functionally differentiated rather than homogeneous. Extending the systems framework identified in adults ^22^, reappraisal-related brain activity in youth was successfully decomposed into reappraisal-specific, common-appraisal, modifiable-emotion and non-modifiable-emotion components, each distributed across conventional network boundaries. This dissociation is consistent with our hypothesis that reappraisal recruits distinct neural components rather than a unitary control system. A region’s functional contribution to reappraisal should depend on its response profile across reappraisal and emotional-reactivity conditions ^22,34^. Task-defined functional organization may thus provide a more informative basis for interpreting distributed neural responses and also corresponding age-related variation than anatomical location or canonical network membership alone ^22,35^. Consistent with this view, the four components showed distinct associations with regulatory success. Reappraisal-specific activity showed the strongest positive association with successful regulation, whereas the emotion-related components showed different and even reverse behavioral relationships. Effective reappraisal therefore appears to depend not only on processes preferentially recruited for regulation, but also on how emotion-related processing is generated and evaluated during reappraisal-like regulation ^35,36^.

The anterior middle temporal gyrus (aMTG) and anterior superior temporal gyrus (aSTG) provide a regional example of this multicomponent organization. The aMTG fell within the reappraisal-specific component, whereas the anatomically adjacent aSTG fell within the common-appraisal component. Activity in both regions statistically accounted for part of the cross-sectional association between age and reappraisal success. Thus, age-related variation in regulatory success intersected multiple functional components rather than a single regulation-selective system, consistent with our hypothesis of a functionally differentiated reappraisal architecture. The localization of both age effects to anterior temporal cortex is also noteworthy. Previous fMRI work has implicated roles of anterior temporal regions in affective meaning during reappraisal, meanwhile independent studies have also linked the anterior temporal lobe to conceptual representations of people and socially relevant knowledge ^37–39^. These findings connect the age-related neural effects to the meaning-reconstruction and social-representational processes highlighted in our multicomponent account of reappraisal ^31,32^. They raise the possibility that age-related differences in reappraisal involve variation in how socially and affectively meaningful information is represented and reframed, alongside variation in cognitive control ^32,40^.

Complementing this functional heterogeneity, multivariate modelling showed that distributed reappraisal responses could be integrated into a generalizable whole-brain neural signature that distinguished reappraisal from passive negative viewing. Reliable discriminative features involved the dorsolateral prefrontal cortex, temporal regions, temporoparietal junction and precuneus, indicating that the generalizable representation was distributed rather than dominated by prefrontal control regions. This pattern is consistent with established models in which reappraisal recruits prefrontal control systems together with temporal and posterior regions involved in representing and reinterpreting the meaning of emotional information delivered by facial expressions ^14,16^. Recent multivariate work likewise shows that reappraisal can be identified from distributed neural patterns that generalize across independent samples and task contexts ^28^. Notably, chronological age explained only a small proportion of individual variation in signature expression. This indicates yet that the distributed neural representation of reappraisal was already broadly reproducible across sampled age range, and age-related variation was more pronounced in the regional and functional organization of this architecture than in its overall multivariate expression ^41,42^. This coexistence of a reproducible whole-brain representation and age-related functional variation further supports our hypothesis that reappraisal develops through reorganization within a distributed architecture rather than uniform strengthening of a focal control system. It also raises a complementary question: whether the interindividual organization of this neural architecture, and its correspondence with regulatory behavior, changes with age.

Intersubject analyses addressed our main concern that age-related differences in reappraisal may be expressed not only in mean responses, but also in how individual differences are organized. Behavioral and neural responses became increasingly similar with age, extending previous evidence that neural representations of emotion become more convergent across individuals from childhood to adolescence ^23^. Importantly, this convergence was stronger during active reappraisal than during passive negative viewing, suggesting that the age-related alignment was particularly associated with regulation rather than reflecting a general convergence of emotional responses. More importantly, the correspondence between neural and behavioral similarity itself varied with age, highlighting that the behavioral relevance of neural variation was not constant across youth. This perspective complements conventional analyses based on developmental group means by treating the structure of interindividual variation, and its brain-behavior mapping, as developmentally informative dimensions ^41,43^. This interpretation is also consistent with recent longitudinal evidence that brain-cognition relationships can become more strongly aligned across early adolescence ^44^. Together, these findings indicate that age-related differences in reappraisal are expressed not only in distributed neural responses, but also in how neural and behavioral differences are organized across individuals and how closely these dimensions correspond.

Several limitations warrant consideration. First, the cross-sectional design captures age-related differences between individuals rather than within-person developmental change, and the interval from 6 to 10.5 years should therefore be interpreted as a period of reliable age-related variation, not as a sensitive developmental period. Second, the four functional components and the aMTG/aSTG findings differ in inferential status, with the latter identifying candidate links rather than causal mechanisms. Third, chronological age subsumes correlated cognitive, pubertal and social processes that cannot be disentangled here. Finally, whether these findings generalize beyond instructed reappraisal of peer-related emotional faces to self-generated strategies and more naturalistic contexts remains to be determined.

In conclusion, emotion regulation via cognitive reappraisal across youth showed coordinated age-related variation across behavioral, functional and interindividual levels. Reappraisal success varied nonlinearly with age. And age-related neural differences were distributed across functionally distinct regulatory, appraisal and emotion-related components. Moreover, the organization of behavioral and neural individual differences also shifted across youth. Against this developmental variation, a distributed neural representation of reappraisal generalized across sites and was detectable throughout the sampled age range. Together, these findings suggest that age-related differences in cognitive reappraisal are better characterized by multilevel reorganization within a distributed, functionally differentiated neural architecture than by uniform strengthening of a focal control system. This framework links when regulatory ability varies with age, how distinct neural components contribute to regulation, and how neural variation maps onto behavior across individuals, providing an integrated account of emotion regulation across youth.

## Methods

### Participants

Participants included in the present study were sourced from the Chinese Child Brain Development program (CCBD), in which the sample was largely recruited via elementary, junior and high schools distributed over east, west, south, north and middle areas of China. The epidemiologically informed sampling procedures were adopted in CCBD to ensure the nationally/populational demographic variation in its cross-sectional sample ranging from 6- to 18-year-olds children and adolescences. From 8 recruitment sites distributed over the whole nation, the current students were invited to participate in the program after the cautious screening for eligibility. The children and adolescences were excluded from the subsequent behavioral tests and brain scanning once meeting one of the exclusion criteria: (1) the presence of contraindications for participating MRI scanning (e.g. implanted metallic device like cardiac pacemakers, claustrophobia etc.); (2) a history of neurological or mental disorder; (3) uncorrected vision, hearing or sensorimotor impairments; (4) inability to understand the instructions or to complete the assessments; (5) unwillingness to complete the assessments. The formally enrolled participants were informed of the details of the assessment contents and procedures, follow which the consent from parents and assent from participants were obtained. The whole protocol of CCBD assessment was approved by appropriate Ethics Review Committee at each recruitment site. And totally 1316 participants entered behavioral analysis.

Participants’ data was included in analyses of the current study if they (1) had completed two runs of the fMRI tasks; (2) effectively controlled their head movements during both scans (mean FD < 0.9 mm); (3) satisfied the least criteria of behavioral performance; (4) had information for covariates of our interests (i.e. age, sex, parental education). Finally, above procedures resulted in 1225 participants included in the following neuroimaging analyses. To our knowledge, the sample size of the current study exceeds those reported in the previously published similar research, which ensures the better statistical power, better ability to detect true effect, and enhanced reproducibility. The whole procedure for screening eligible participants can refer to *Supplementary Figure 1*.

### fMRI task and stimuli

As depicted in **Fig. 1a**, the emotion regulation paradigm starts with a star fixation of duration in a range of 1-5 s, and then a picture representing the target agent appears with a negative (fear, disgust, anger and sadness) or neutral facial expression, under which a sign informs participants of whether using cognitive reappraisal to down-regulate the self-experience of negative emotions towards target (sign: downward arrow) or just passive looking at the picture (sign: eye). Subsequently in the next screen lasting for 6 s, the participant needs to evaluate their real-time emotional feelings on a 4-point Likert scale spanning from ‘very good’ to ‘very bad’. Given the complexity of our task, the instructions were presented to the participants in the video form, which offered a detailed and vivid description regarding the cognitive reappraisal as well as the whole experimental procedure. This facilitated the understanding of the task, and also avoided the bias introduced by the inconsistency of introduction processes.

### fMRI data acquisition and pre-processing

T1-weighted structural and task-based functional MRI data were acquired across Siemens 3T Prisma, UH uMR890, and GE MR750 scanners. Preprocessing was performed using the typical fMRIPrep workflow ^45^. The first 10 volumes (7.5 s) of each functional run were discarded to allow for magnetization equilibrium and participant habituation. Functional images underwent head motion correction via rigid-body registration to a single-band reference image, and susceptibility-induced distortion correction using fieldmaps. The distortion-corrected BOLD reference was co-registered to the skull-stripped T1-weighted structural image using boundary-based registration and subsequently normalized to MNI152NLin2009cAsym space. To minimize spatial interpolation artifacts, all motion, co-registration, distortion correction, and standard space transformation matrices were concatenated and applied to the BOLD time series in a single step.

Functional runs were retained for analysis if at least 80% of the volumes exhibited a framewise displacement (FD) < 0.9 mm ^46^. To mitigate physiological and motion artifacts, nuisance signals were regressed out using a General Linear Model (GLM). Nuisance regressors included the global signal, mean white matter and cerebrospinal fluid signals, CompCor components, FD, head motion parameters (either standard 6 or Friston-24), and linear/quadratic temporal trends; the model’s constant term was preserved to maintain image-to-background contrast. Finally, low-frequency temporal drift was removed via high-pass filtering (> 0.0078 Hz) with the temporal mean added back, and data were spatially smoothed using a full-width-at-half-maximum (FWHM) Gaussian kernel of 6 mm.

### First-level analysis

Within-subject task effects and contrasts were estimated with a general linear model (GLM) using SPM12 ^47^. The three emotion-regulation conditions (i.e. reappraisal of negative faces (Reappraise), passive viewing of negative faces (Look), and passive viewing of neutral faces (Neutral)) were separately modeled with boxcar functions and convolved with the canonical hemodynamic response function (HRF). Regressors included one for the period of picture presentation in each task condition. An additional zero-duration regressor modeled motor effects on trials with a recorded response. The timing and duration of stimulus epochs were modeled according to their recorded onset and duration and aligned to the onset of the first fixation period. A high-pass filter with a 128-s cutoff was applied to remove low-frequency temporal drift. Nuisance regressors included the Friston 24-parameter motion model ^48^, comprising the six rigid-body motion parameters, their temporal derivatives, and the squared terms of both; the first six anatomical CompCor components; and separate indicator regressors for volumes identified during preprocessing as motion outliers or non-steady-state volumes. Temporal autocorrelation was modeled using a first-order autoregressive [AR(1)] process. Subject level contrasts of interest included cognitive reappraisal defined by Reappraise - Look and emotion reactivity defined by Look – Neutral.

### Generalized additive modeling of age-related developmental changes

Developmental trajectories were modeled using generalized additive models (GAMs). For repeated-measures outcomes, subject-level within-participant contrasts (Reappraise - Look) were first calculated and analyzed separately, yielding one observation per participant for each contrast. Age was entered as a penalized smooth term. When sex-specific trajectories were examined, models additionally included a sex factor and a sex-specific smooth deviation:

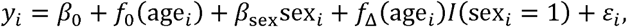

where *f*_0_(age) represents the developmental trajectory of the reference sex and *f*_Δ_(age) represents the age-dependent deviation of the other sex. Smoothing parameters were selected by grid search. Age windows of significant age-related change were determined by estimating the first derivative of the age smooth function via finite differences. Statistical significance was established when the simultaneous 95% confidence interval for this derivative did not contain zero ^49^.

As indicated Fig. 2a and consistent with Mousley et al. (2025) ^50^, we also identified the turning points in developmental trajectories. To characterize uncertainty in trajectory shape, coefficient vectors were repeatedly sampled from a multivariate normal approximation: 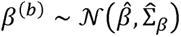, where 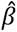 and 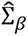 denote the fitted GAM coefficients and their estimated covariance matrix. Each draw generated a posterior trajectory *f*^(*b*)^(*a*), from which the first derivative

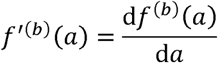

was calculated. Candidate turning points of trajectory were defined as ages at which the derivative changed sign, with positive-to-negative transitions classified as peaks and negative-to-positive transitions as valleys. Candidates near the boundaries of the observed age range were excluded. And stable turning points were identified from the distribution of posterior candidate ages using kernel density estimation (KDE):

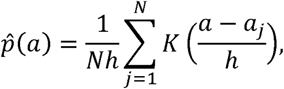

where *a_j_* denotes a posterior candidate turning-point age, *K* is the kernel function, and *h* is the bandwidth. Local KDE modes were treated as candidate final turning points, and posterior candidates within a predefined age window around each mode were assigned to that turning point. Final turning points were retained only when they exceeded prespecified thresholds for the number and proportion of posterior draws supporting a local sign change.

#### Generalized additive modeling of age-related effect on brain response of emotion regulation

GAM was also used to characterize nonlinear age-related variation in regional brain responses associated with emotion regulation, following the general analytic framework presented in Sydnor et al. (2023) ^51^. For each participant, the subject-level emotion-regulation contrast map (i.e. Reappraise - Look) was summarized within each cortical parcel by averaging values across voxels ^24^, yielding one regional contrast estimate per participant. Separate GAMs were then fitted for each cortical parcel, with the regional emotion-regulation response as the dependent variable, age modeled as a smooth function, and prespecified covariates entered as linear terms including sex, mean FD, site ids and degree of parental education.

The age effect was modeled using penalized spline basis functions with a maximum basis complexity of (k=3). For each regional model, the smoothing penalty was selected by generalized cross-validation over a logarithmically spaced candidate grid. Statistical significance of the age effect was evaluated by comparing the full GAM with an otherwise identical reduced model that omitted the smooth age term. Model improvement was quantified from the reduction in residual sum of squares while accounting for the effective degrees of freedom of the full and reduced models. Resulting *p* values were corrected across cortical parcels using the Benjamini–Hochberg false discovery rate procedure (FDR *q* < 0.05). And the magnitude of the age effect was quantified using partial R^2^, defined by the improvement in model fit between the full model containing the smooth age term and the corresponding reduced model without age term.

#### Reappraisal-specific and common brain systems using Bayesian identification approach

Adapted from Bo et al. (2024) ^22^, as Fig. 4a indicated, we used an axiomatic systems-identification approach to identify four brain components according to their joint response profiles across emotion regulation and emotion reactivity. For each participant, contrast images were computed for emotion regulation (Reappraise - Look) and emotion reactivity (Look - Neutral). Each contrast was entered separately into a mass-univariate second-level model, and the resulting voxel-wise t statistics were converted to Bayes factors using the Jeffreys–Zellner–Siow (JZS) prior ^52^:

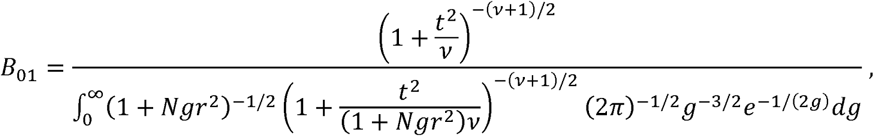

Where *N* denotes sample size, *t* for the voxel-wise group-level t statistic, *v* for the degrees of freedom, and *r* for the prior scale parameter set to 0.707. Consistent with previous studies ^22,53^, BF_10_ > 10 was considered strong evidence for an effect and BF_10_ < 0.1 as strong evidence for the null hypothesis.

Thresholded Bayes-factor and effect-direction maps were then combined by logical conjunction. For voxel *v*, affiliation with each system was defined as:

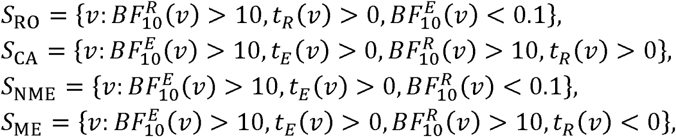

where *R* and *E* denote the emotion regulation and emotion reactivity contrasts, respectively. These criteria identified reappraisal-only (RO), common-appraisal (CA), non-modifiable emotion-generation (NME), and modifiable emotion-generation (ME) functional components. Reappraisal-only voxels were additionally required to show positive mean activation during reappraising negative faces, whereas voxels in the other three systems were required to show positive mean activation during reactivity to negative faces. The resulting conjunction maps were converted to binary masks, and clusters smaller than 15 contiguous voxels were removed.

Subsequently, we identified potential age-related regions within functional components through conjunction between brain mask of GAM-derived age-related regions and brain masks of four brain components (Fig. 4d). Then these conjunctive regions were utilized to build mediation models, in which brain activation in these regions was considered as mediators to link age variations across youth and individual differences in reappraisal success.

#### Multivariate modeling for neural signature of emotion regulation

Parcel-wise task-evoked beta maps were used as features for multivariate pattern analysis ^24^. Each participant contributed one map for cognitive reappraisal of negative stimuli (Reappraise) and one map for passive viewing of negative stimuli (Look). Reappraise was coded as the positive class (1) and Look as the reference class (0). Subject and acquisition-site identifiers were retained throughout the analysis. Three classification models (logistic L2 regression, linear support vector machine and radial-basis-function support vector machine) were simultaneously used to identify a distributed neural pattern discriminating Reappraise from Look in youth. Model performance was evaluated using nested leave-one-site-out cross-validation. In the outer loop, one acquisition site was held out as an independent test set and the remaining sites were used for model development. Within each outer training set, hyperparameters were selected using leave-one-site-out cross-validation across the remaining training sites. Feature standardization and all model-fitting steps were performed within the training data only. The selected model was then refitted to the complete outer training set and applied to the held-out site. Model performance was summarized using accuracy, balanced accuracy, sensitivity, specificity, ROC curves, and area under the ROC curve. Overall performance was calculated from the pooled out-of-fold predictions, and fold-specific results were additionally examined to assess cross-site consistency.

Statistical significance of predictive performance was assessed using within-subject label permutation. For each permutation, the Reappraise and Look labels were randomly exchanged within participants, preserving the paired design, site structure, and class balance. The complete nested cross-validation procedure was repeated for each permuted dataset to generate a null distribution of model performance. Empirical *p* values were calculated as the proportion of permuted performance values equal to or greater than the observed value. The stability of the final multivariate pattern (i.e. decoder weights) was assessed using subject-level bootstrap resampling. Participants were sampled with replacement while retaining both condition-specific maps from each selected participant. The classifier was refitted in each bootstrap sample using the final selected hyperparameters, and the resulting pattern was retained. For each feature, a bootstrap ratio was calculated as the mean bootstrap value divided by its bootstrap standard deviation and was interpreted as an index of contribution magnitude and resampling stability. Directional stability was additionally assessed from the proportion of bootstrap estimates with opposite signs. Two-sided sign-stability *p* values were corrected for multiple comparisons using the FDR procedure at *q* < 0.05.

Since linear decoding weights are not directly interpretable as regional activation effects, they were transformed into reconstructed activation patterns using the Haufe transformation ^27^. The transformation combined the covariance structure of the training features with the fitted decoding weights and model scores, yielding feature values that reflect the direction of association between regional activation and the latent process captured by the classifier. Positive reconstructed activation values indicated greater expression of the reappraisal-related process, whereas negative values indicated the relative expression of the passive viewing negative condition. Haufe patterns were calculated for the final model and for each bootstrap sample, allowing both bootstrap-ratio and FDR-based stability maps to be derived. To demonstrate spatial correspondence between multivariate pattern (i.e. neural signature) and univariate activation, we also computed the spatial similarities between Haufe transformed pattern and brain activation of cognitive reappraisal and emotion reactivity. The spatial similarity with canonical brain networks ^25^ was also computed using cosine similarity to address network contributions to reappraisal neural signature.

Subsequently, the signature response was defined as the continuous out-of-fold classifier score. Within-subject Reappraise-minus-Look differences of the signature response were subsequently used in individual-difference analyses for age effect, in which both GAM and GLM were conducted. The whole procedure for this section of analysis can be seen in Fig. 5a.

#### Inter-subject similarity analysis for developmental reorganization involved in reappraisal

We tested whether individual differences in neural and behavioral emotion representations were organized according to alternative models of development using an inter-subject representational similarity analysis framework.

Single-trial activation maps at reappraisal and passive viewing conditions were estimated using a least-squares-separate (LSS) general linear model. A separate first-level model was fitted for each trial, with the target trial modeled by an individual regressor and all remaining trials combined into a separate nuisance regressor. Event regressors were convolved with the SPM canonical hemodynamic response function and its temporal derivative, with cosine drift terms, high-pass filtering, and an AR(1) noise model included in model estimation. The beta coefficient corresponding to the target trial was retained, yielding one activation map for each trial and participant. Within each of 232 predefined regions of interest ^24,54^, voxel-wise beta estimates were extracted to form trial-specific multivoxel activation patterns.

As indicated in **Fig. 6a**, trials were grouped into four emotion categories, i.e. anger, fear, disgust, and sadness, and multivoxel activation patterns were averaged within each category to obtain an emotion-specific prototype pattern for each participant and ROI. For emotion e, the prototype pattern was defined as:

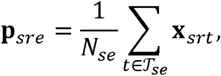

where **x***_srt_* denotes the multivoxel pattern for participant *s*, ROI *r*, and trial *t*, and T*_se_* denotes the set of trials belonging to emotion *e*. For each participant and ROI, the four emotion prototype patterns were cocktail-blank centered by removing, for each voxel, its mean activation across the four emotion categories. Pairwise similarity between the centered prototype patterns was then calculated using Spearman correlation, producing a 4 × 4 emotion-by-emotion representational similarity matrix (RSM). Because the RSM was symmetric, its six unique off-diagonal upper-triangular elements were extracted to form a participant-specific representational profile. These correlation coefficients were Fisher r-to-z transformed before cross-participant comparison. Thus, each participant was represented by a six-dimensional vector describing the relative organization of the four emotion categories within a given ROI.

Then, representational dissimilarity between every pair of participants was quantified using Mahalanobis distance, which accounts for covariance among the six representational dimensions:

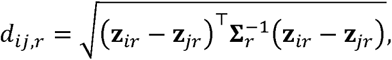

Where **z***_ir_* and **z***_jr_* denote the Fisher-transformed representational profiles of participants *i* and *j*, respectively, and **Σ***_r_* represents the covariance structure across representational dimensions. Covariance estimation used ridge regularization with a pseudoinverse as the default implementation (ridge parameter = 10⁻⁶). Mahalanobis distances were subsequently sign-reversed and standardized across off-diagonal participant pairs so that larger values indicated greater representational similarity; diagonal elements were set to 1. This procedure yielded one symmetric N*N ISS matrix for each ROI and experimental condition, with each off-diagonal element indexing the degree to which a given pair of participants shared a similar organization of neural representations. Additionally, behavioral inter-subject similarity was quantified from participants’ trial-wise emotion self-ratings separately for the reappraisal and passive viewing conditions. For each participant, trials were grouped into four emotion categories, i.e. anger, fear, disgust, and sadness, and self-ratings were averaged across trials within each category, yielding a four-dimensional emotion-rating profile representing each participant’s pattern of subjective emotional responses across the four emotion categories. Pairwise dissimilarity between participants was then quantified using Mahalanobis distance between their emotion-rating profiles, which was followed by sign-reversion and standardization of distances across off-diagonal participant pairs. These pairwise ISS matrices were used in all subsequent developmental and brain-behavior similarity analyses.

Following previous studies ^23^, three developmental model were constructed from chronological age: nearest-neighbor model using *a*_max_−∣ *a_i_* − *a_j_* ∣; convergence model using min (*a_i_*, *a_j_*); and divergence model using *a*_max_ − (*a_i_* + *a_j_*)/2. The fit of each developmental model was quantified as the Spearman correlation between the vectorized upper triangles of the developmental and observed ISS matrices. Significance was assessed using 10,000 Mantel-style permutations in which participant labels were randomly permuted simultaneously across rows and columns ^43,55^. This procedure was repeated separately for behavioral ISS and for each region-specific brain ISS matrix in each condition. To further characterize condition-related changes in inter-subject similarity, difference matrices between conditions were calculated as ΔISS = ISS_Reappraise_ – ISS_Look_. For candidate regions surviving the exploratory uncorrected *p* < 0.05 developmental-model screening threshold, permutation-based matrix regression was used to predict Δbehavioral ISS from Δbrain ISS, the developmental model matrices, and their interactions (lower panel of Fig. 6a). Continuous predictors were standardized before interaction terms were calculated. Statistical significance of regression coefficients was determined using 10,000 simultaneous row-column permutations of participant labels, thereby preserving the dependency structure of the dyadic observations.

## Data & code availability

The data that support the findings of this study are available from the corresponding authors upon reasonable request, subject to applicable ethical and institutional data-sharing restrictions.

## Supporting information

supplementary informations

## Acknowledgements

This work was supported by the Brain Science and Brain-like Intelligence Technology-National Science and Technology Major Project (2021ZD0200500).

## Author contributions

R.D., Z.Ze. and S.Q. conceived and designed the study. R.D. and Z.Zh. performed the analyses and prepared the figures. R.Q., B.H., Q.H., Y.L., Y.H. and X.Z. contributed to data collection, data curation and interpretation of the results. Z.Ze. drafted the manuscript. S.Q. supervised the study. All authors reviewed and approved the final manuscript.

