## supplementary informations for "Functional reorganization of large-scale brain systems underlies emotion regulation maturation"

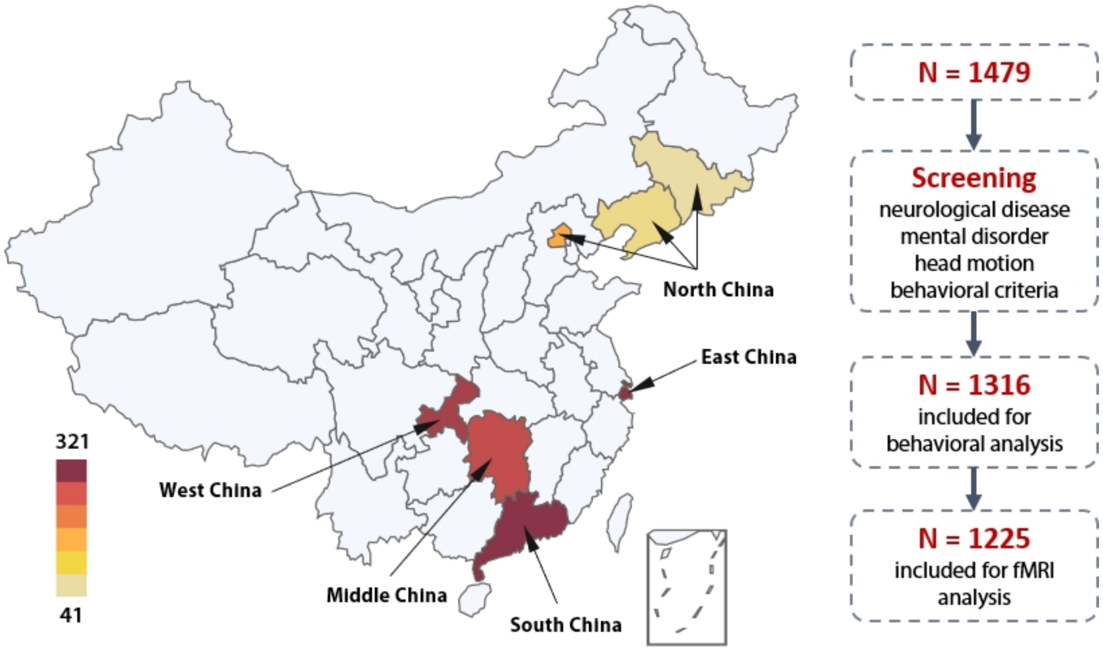


**Supplementary Figure 1.** The distribution of scanning sites in China and screening procedure for behavioral and fMRI analysis.

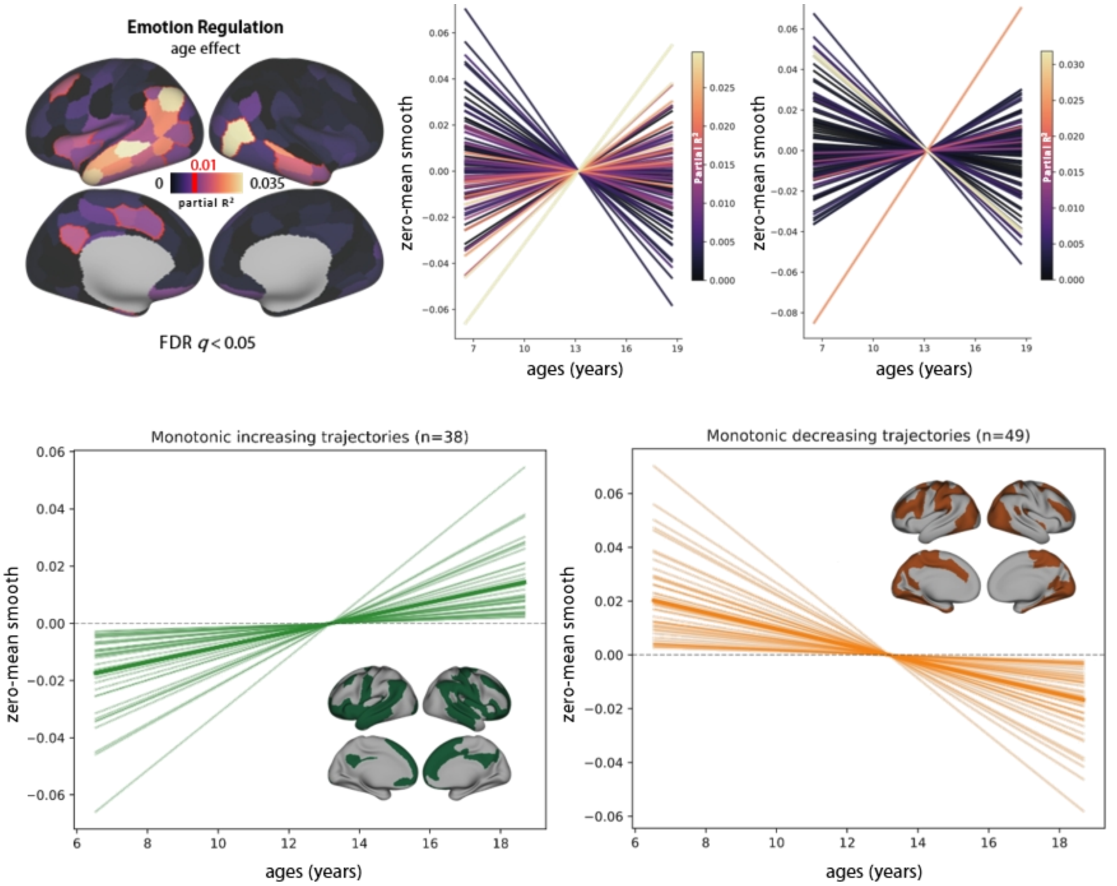


**Supplementary Figure 2.** The age-dependent trajectories of emotion regulation and groups of brain regions/parcels that demonstrate monotonically increasing (green) and decreasing (orange) developmental trends with age.

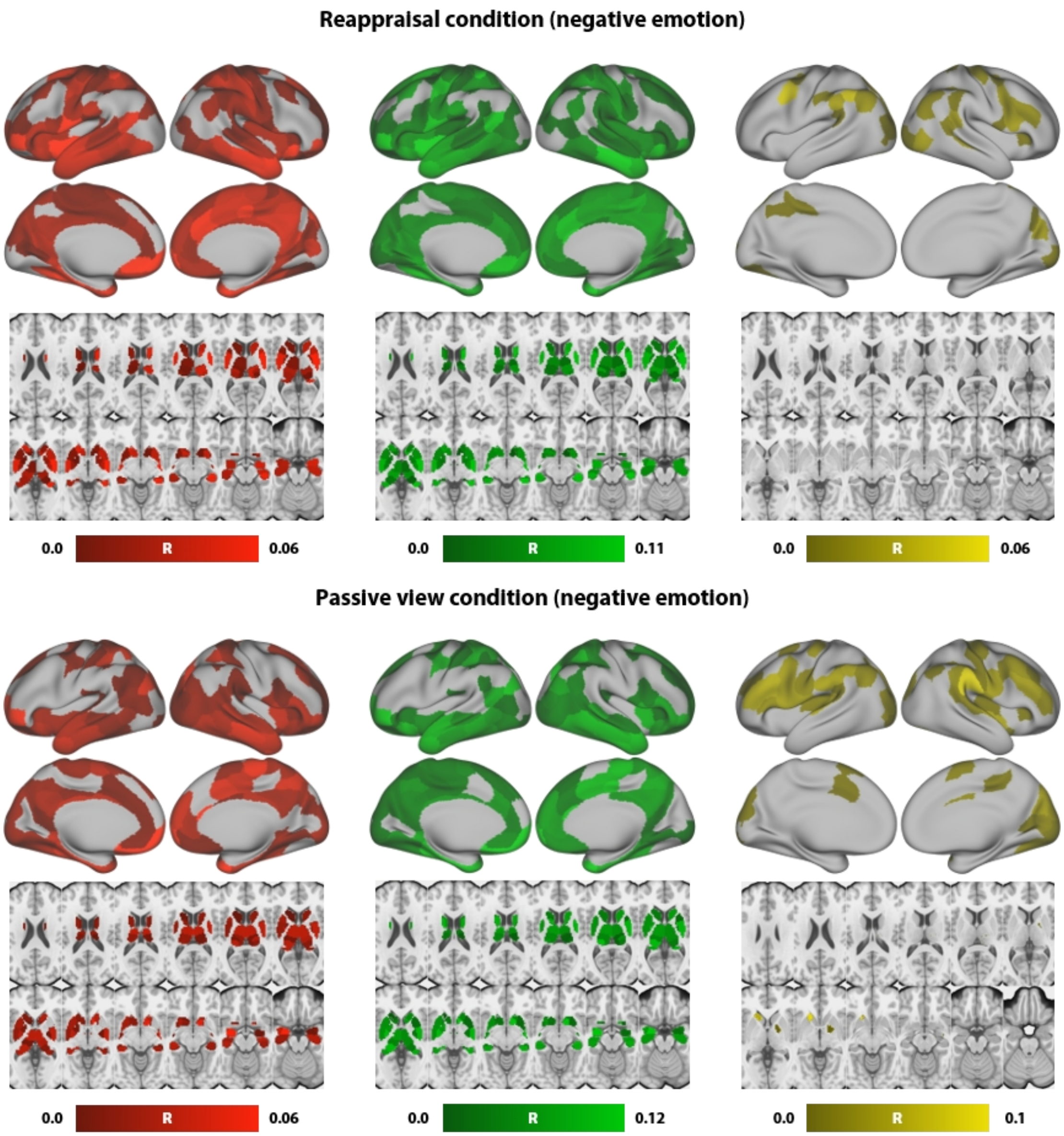


**Supplementary Figure 3.** The brain maps of associations between age/developmental model and inter-individual differences of brain activity during emotion regulation. Here we only visualized the positive associations (R), since the negative association is meaningless.

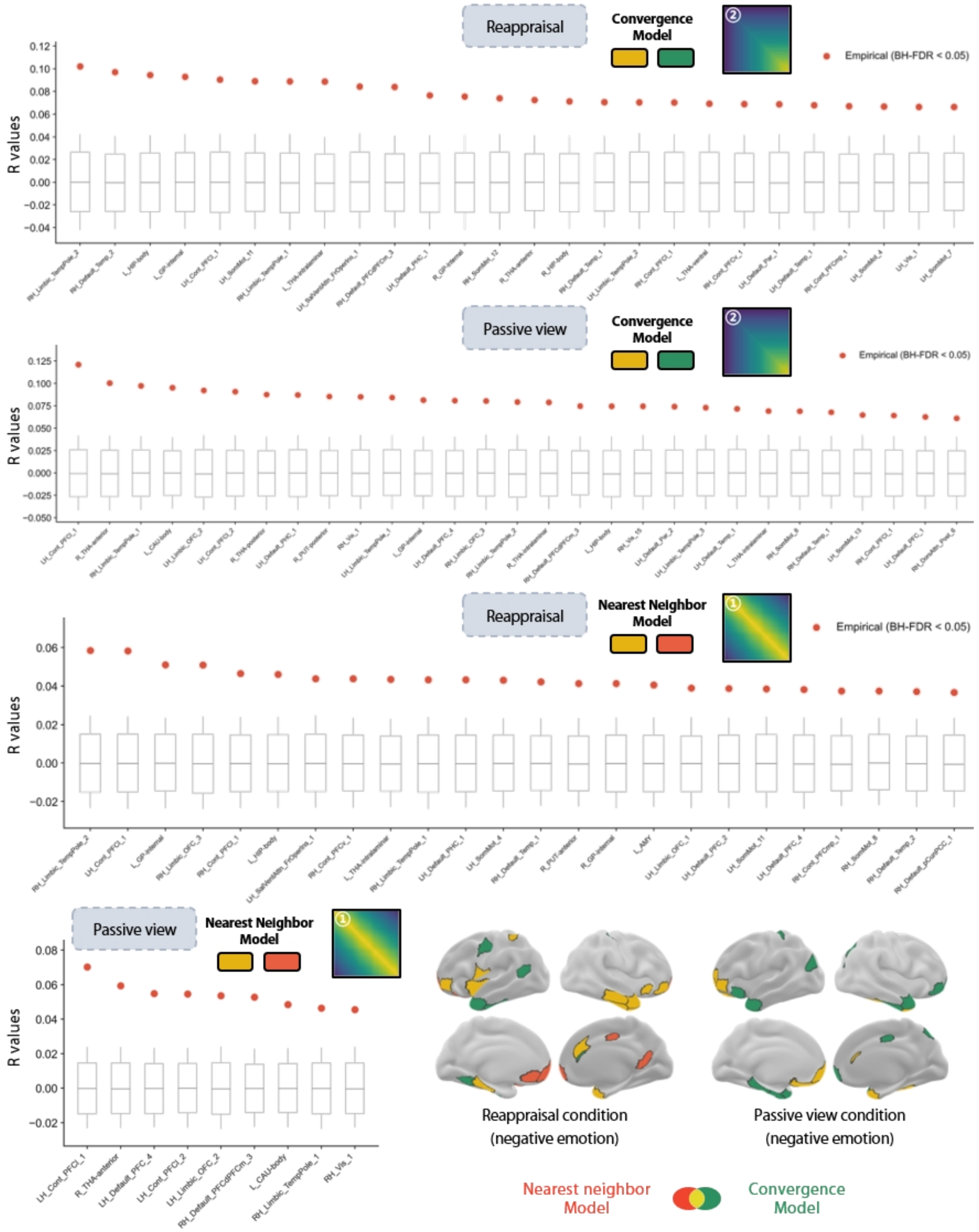


**Supplementary Figure 4.** The brain regions/parcels surviving multiple comparisons correction, and boxplots showing statistical results of multiple correction comparisons (FDR $p < 0.05$), in which (1) red points represent observed R values of associations between inter-individual differences of brain activations and age/developmental models, and (2) boxes along with whiskers represent 0% - 100% percentiles of permuted R values of inter-individual differences of brain activations and age/developmental models.

**Supplementary Table 1.** The brain regions significantly activated during emotion regulation (Reappraise - Look at FDR $q < 0.005$ with minimum $k = 30$)

| Cluster | Hemi. | MNI coordinates x | y | z | Peak stats | Volume ($mm^3$) | Cluster overlaps with AAL |
|---|---|---|---|---|---|---|---|
| **1** | **L** | **-52.1** | **-59.3** | **25.9** | **13.86** | **91141** | 36.37% Temporal_Mid_L; 16.53% no_label; 9.62% Temporal_Inf_L; 8.58% Angular_L; 5.01% Occipital_Mid_L |
| | L | -42.5 | -56.9 | 25.9 | 12.70 | | |
| | L | -61.7 | -37.7 | -0.5 | 12.50 | | |
| | L | -54.5 | 0.7 | -24.5 | 12.39 | | |
| **2** | **R** | **51.1** | **-73.7** | **1.9** | **13.59** | **51480** | 49.41% Temporal_Mid_R; 9.96% Temporal_Inf_R; 8.75% Angular_R; 8.11% Temporal_Sup_R; 7.22% Occipital_Mid_R; 6.87% Temporal_Pole_Mid_R |
| | R | 60.7 | -40.1 | -0.5 | 8.91 | | |
| | R | 51.1 | 0.7 | -24.5 | 8.80 | | |
| | R | 51.1 | -54.5 | 25.9 | 8.78 | | |
| **3** | **L** | **-8.9** | **-47.3** | **35.5** | **8.98** | **12427** | 50.61% Precuneus_L; 13.68% Cingulate_Post_L; 10.68% Precuneus_R; 9.57% Cingulate_Mid_L; 6.12% Cuneus_L |
| | L | -4.1 | -64.1 | 28.3 | 6.63 | | |
| | R | 7.9 | -49.7 | 33.1 | 5.29 | | |
| | R | 10.3 | -56.9 | 40.3 | 4.85 | | |
| **4** | **L** | **-44.9** | **10.3** | **52.3** | **8.40** | **11363** | 57.18% Frontal_Mid_2_L; 28.71% Frontal_Sup_2_L; 13.50% Precentral_L |
| | L | -16.1 | 41.5 | 49.9 | 5.73 | | |
| | L | -20.9 | 22.3 | 54.7 | 5.38 | | |
| | L | -16.1 | 43.9 | 42.7 | 5.30 | | |
| **5** | **R** | **15.1** | **-85.7** | **-41.3** | **8.34** | **2128** | 87.01% Cerebellum_Crus2_R; 12.99% no_label |
| **6** | **R** | **24.7** | **-83.3** | **21.1** | **6.84** | **3801** | 63.64% Occipital_Sup_R; 34.91% Occipital_Mid_R |
| **7** | **R** | **27.1** | **-64.1** | **-5.3** | **6.74** | **4161** | 60.80% Lingual_R; 14.29% Cerebellum_Crus1_R; 12.29% Fusiform_R; 5.98% Cerebellum_6_R |
| | R | 17.5 | -85.7 | -17.3 | 6.04 | | |
| | R | 10.3 | -83.3 | -12.5 | 5.37 | | |
| | R | 17.5 | -76.1 | -12.5 | 4.74 | | |

| Cluster | Hemi. | MNI coordinates | | | Peak stats | Volume (mm$^3$) | Cluster overlaps with AAL |
|---|---|---|---|---|---|---|---|
| **8** | **L** | **-23.3** | **-85.7** | **-38.9** | **5.92** | **1327** | 89.58% Cerebellum_Crus2_L; 10.42% no_label |
| **9** | **L** | **-4.1** | **55.9** | **-10.1** | **5.90** | **1410** | 75.49% Frontal_Med_Orb_L; 18.63% Rectus_L; 5.88% Frontal_Med_Orb_Ra |
| **10** | **R** | **12.7** | **43.9** | **47.5** | **5.64** | **2529** | 61.20% Frontal_Sup_2_R; 38.80% Frontal_Sup_Medial_R |
| | R | 15.1 | 51.1 | 37.9 | 5.33 | | |
| | R | 12.7 | 36.7 | 49.9 | 5.08 | | |
| **11** | **R** | **46.3** | **29.5** | **-14.9** | **5.06** | **746** | 53.70% Frontal_Inf_Orb_2_R; 22.22% OFCpost_R; 20.37% OFClat_R |
| | R | 51.1 | 34.3 | -2.9 | 3.94 | | |

**Supplementary Table 2.** The brain regions significantly activated during emotion reactivity ( Look - Neutral at FDR $q < 0.005$ with minimum $k = 30$)

| Cluster | Hemi. | MNI coordinates | | | Peak stats | Volume ($mm^3$) | Cluster overlaps with AAL |
|---|---|---|---|---|---|---|---|
| | | x | y | z | | | |
| **1** | **R** | **48.7** | **-37.7** | **6.7** | **18.62** | 97334 | 13.08% Temporal_Mid_R; 9.98% no_label; 8.62% Temporal_Sup_R; 7.44% Frontal_Inf_Tri_R; 7.30% Temporal_Pole_Sup_R; 6.97% Temporal_Inf_R; 6.68% Fusiform_R; 5.58% Temporal_Pole_Mid_R; 5.30% Insula_R; 5.24% Occipital_Inf_R |
| | R | 51.1 | 24.7 | -2.9 | 15.89 | | |
| | R | 51.1 | 12.7 | -24.5 | 15.29 | | |
| | R | 51.1 | -23.3 | -2.9 | 15.23 | | |
| **2** | **L** | **-44.9** | **24.7** | **-5.3** | **16.16** | 162805 | 13.16% no_label; 9.21% Temporal_Mid_L; 8.34% Frontal_Sup_Medial_L; 7.01% Frontal_Inf_Tri_L; 5.27% Temporal_Pole_Sup_L; 5.00% Frontal_Sup_2_L |
| | L | -42.5 | 19.9 | -12.5 | 15.03 | | |
| | L | -30.5 | -90.5 | -10.1 | 14.85 | | |
| | L | -6.5 | 53.5 | 33.1 | 13.45 | | |
| **3** | **L** | **-11.3** | **-71.3** | **35.5** | **6.62** | 1479 | 63.55% Precuneus_L; 26.17% Cuneus_L; 5.61% Occipital_Sup_L |
| **4** | **R** | **22.3** | **-80.9** | **-31.7** | **6.21** | 1244 | 60.00% Cerebelum_Crus1_R; 40.00% Cerebelum_Crus2_R |
| **5** | **R** | **7.9** | **10.3** | **6.7** | **5.81** | 2032 | 59.18% Caudate_R; 40.82% no_label |
| | R | 10.3 | 3.1 | 11.5 | 4.66 | | |
| | R | 10.3 | -1.7 | 18.7 | 4.60 | | |
| | R | 7.9 | 0.7 | 4.3 | 3.95 | | |
| **6** | **L** | **-4.1** | **-25.7** | **28.3** | **5.68** | 2308 | 65.27% no_label; 17.37% Cingulate_Mid_L; 8.98% Cingulate_Post_L; 8.38% Cingulate_Mid_R |
| | R | 5.5 | -20.9 | 28.3 | 5.36 | | |
| | L | -4.1 | -40.1 | 23.5 | 4.07 | | |
| | L | -4.1 | -23.3 | 40.3 | 3.94 | | |
| **7** | **L** | **-28.1** | **-68.9** | **-50.9** | **5.42** | 594 | 60.47% Cerebelum_8_L; 20.93% no_label; 18.60% Cerebelum_7b_L |

| Cluster | Hemi. | MNI coordinates | | | Peak stats | Volume (mm$^3$) | Cluster overlaps with AAL |
|---|---|---|---|---|---|---|---|
| **8** | **L** | **-4.1** | **-56.9** | **-36.5** | **5.40** | 2128 | 42.21% Vermis_9; 17.53% Cerebelum_9_R; 14.29% Vermis_8; 9.74% Cerebelum_9_L; 6.49% Cerebelum_8_L; 5.19% no_label |
| | R | 7.9 | -61.7 | -43.7 | 5.02 | | |
| **9** | **L** | **-18.5** | **-78.5** | **-34.1** | **5.20** | 1410 | 50.00% Cerebelum_Crus2_L; 26.47% no_label; 20.59% Cerebelum_Crus1_L |
| | L | -13.7 | -80.9 | -43.7 | 5.03 | | |
| **10** | **R** | **3.1** | **53.5** | **-22.1** | **4.26** | 483 | 77.14% Rectus_L; 22.86% Rectus_R |
| | L | -4.1 | 60.7 | -22.1 | 3.73 | | |
| **11** | **L** | **-42.5** | **-25.7** | **54.7** | **4.16** | 1216 | 75.00% Postcentral_L; 25.00% Precentral_L |
| **12** | **R** | **29.5** | **51.1** | **-2.9** | **3.97** | 414 | 43.33% Frontal_Sup_2_R; 40.00% Frontal_Mid_2_R; 16.67% no_label |

**Supplementary Table 3.** The brain regions/parcels in which inter-individual similarity of brain response during emotion regulation and reactivity is significantly aligned with developmental models (FDR $q < 0.05$)

| condition | developmental model | parcel/region labels [a] | Spearman's r |
|---|---|---|---|
| passive viewing (negative) | convergence | LH_Cont_PFCl_1 | 0.121 |
| passive viewing (negative) | convergence | R_THA-anterior | 0.100 |
| passive viewing (negative) | convergence | RH_Limbic_TempPole_1 | 0.097 |
| passive viewing (negative) | convergence | L_CAU-body | 0.095 |
| passive viewing (negative) | convergence | LH_Limbic_OFC_2 | 0.092 |
| passive viewing (negative) | convergence | LH_Cont_PFCl_2 | 0.091 |
| passive viewing (negative) | convergence | R_THA-posterior | 0.087 |
| passive viewing (negative) | convergence | LH_Default_PHC_1 | 0.087 |
| passive viewing (negative) | convergence | R_PUT-posterior | 0.085 |
| passive viewing (negative) | convergence | RH_Vis_1 | 0.085 |
| passive viewing (negative) | convergence | LH_Limbic_TempPole_1 | 0.084 |
| passive viewing (negative) | convergence | L_GP-internal | 0.081 |
| passive viewing (negative) | convergence | LH_Default_PFC_4 | 0.081 |
| passive viewing (negative) | convergence | RH_Limbic_OFC_3 | 0.081 |
| passive viewing (negative) | convergence | RH_Limbic_TempPole_2 | 0.079 |
| passive viewing (negative) | convergence | R_THA-intralaminar | 0.079 |
| passive viewing (negative) | convergence | RH_Default_PFCdPFCm_3 | 0.075 |
| passive viewing (negative) | convergence | L_HIP-body | 0.075 |
| passive viewing (negative) | convergence | RH_Vis_15 | 0.074 |
| passive viewing (negative) | convergence | LH_Default_Par_2 | 0.074 |
| passive viewing (negative) | convergence | LH_Limbic_TempPole_3 | 0.073 |

| | | | |
|---|---|---|---|
| passive viewing (negative) | convergence | LH_Default_Temp_1 | 0.072 |
| passive viewing (negative) | convergence | L_THA-intralaminar | 0.069 |
| passive viewing (negative) | convergence | RH_SomMot_8 | 0.069 |
| passive viewing (negative) | convergence | RH_Default_Temp_1 | 0.068 |
| passive viewing (negative) | convergence | LH_SomMot_13 | 0.065 |
| passive viewing (negative) | convergence | RH_Cont_PFCl_1 | 0.064 |
| passive viewing (negative) | convergence | LH_Default_PFC_1 | 0.063 |
| passive viewing (negative) | convergence | RH_DorsAttn_Post_6 | 0.061 |
| passive viewing (negative) | nearest neighbor | LH_Cont_PFCl_1 | 0.070 |
| passive viewing (negative) | nearest neighbor | R_THA-anterior | 0.059 |
| passive viewing (negative) | nearest neighbor | LH_Default_PFC_4 | 0.055 |
| passive viewing (negative) | nearest neighbor | LH_Cont_PFCl_2 | 0.055 |
| passive viewing (negative) | nearest neighbor | LH_Limbic_OFC_2 | 0.054 |
| passive viewing (negative) | nearest neighbor | RH_Default_PFCdPFCm_3 | 0.053 |
| passive viewing (negative) | nearest neighbor | L_CAU-body | 0.049 |
| passive viewing (negative) | nearest neighbor | RH_Limbic_TempPole_1 | 0.046 |
| passive viewing (negative) | nearest neighbor | RH_Vis_1 | 0.046 |
| Reappraisal (negative) | convergence | RH_Limbic_TempPole_2 | 0.102 |
| Reappraisal (negative) | convergence | RH_Default_Temp_2 | 0.097 |
| Reappraisal (negative) | convergence | L_HIP-body | 0.095 |
| Reappraisal (negative) | convergence | L_GP-internal | 0.093 |
| Reappraisal (negative) | convergence | LH_Cont_PFCl_1 | 0.090 |
| Reappraisal (negative) | convergence | LH_SomMot_11 | 0.089 |
| Reappraisal (negative) | convergence | RH_Limbic_TempPole_1 | 0.089 |
| Reappraisal (negative) | convergence | L_THA-intralaminar | 0.089 |
| Reappraisal (negative) | convergence | LH_SalVentAttn_FrOperIns_1 | 0.084 |
| Reappraisal (negative) | convergence | RH_Default_PFCdPFCm_3 | 0.084 |
| Reappraisal (negative) | convergence | LH_Default_PHC_1 | 0.077 |
| Reappraisal (negative) | convergence | R_GP-internal | 0.076 |
| Reappraisal (negative) | convergence | RH_SomMot_12 | 0.074 |

| | | | |
|---|---|---|---|
| Reappraisal (negative) | convergence | R_THA-anterior | 0.073 |
| Reappraisal (negative) | convergence | R_HIP-body | 0.071 |
| Reappraisal (negative) | convergence | RH_Default_Temp_1 | 0.071 |
| Reappraisal (negative) | convergence | LH_Limbic_TempPole_2 | 0.070 |
| Reappraisal (negative) | convergence | RH_Cont_PFCl_1 | 0.070 |
| Reappraisal (negative) | convergence | L_THA-ventral | 0.069 |
| Reappraisal (negative) | convergence | RH_Cont_PFCv_1 | 0.069 |
| Reappraisal (negative) | convergence | LH_Default_Par_1 | 0.069 |
| Reappraisal (negative) | convergence | LH_Default_Temp_1 | 0.068 |
| Reappraisal (negative) | convergence | RH_Cont_PFCmp_1 | 0.067 |
| Reappraisal (negative) | convergence | LH_SomMot_4 | 0.067 |
| Reappraisal (negative) | convergence | LH_Vis_1 | 0.066 |
| Reappraisal (negative) | convergence | LH_SomMot_7 | 0.066 |
| Reappraisal (negative) | nearest neighbor | RH_Limbic_TempPole_2 | 0.058 |
| Reappraisal (negative) | nearest neighbor | LH_Cont_PFCl_1 | 0.058 |
| Reappraisal (negative) | nearest neighbor | L_GP-internal | 0.051 |
| Reappraisal (negative) | nearest neighbor | RH_Limbic_OFC_3 | 0.051 |
| Reappraisal (negative) | nearest neighbor | RH_Cont_PFCl_1 | 0.047 |
| Reappraisal (negative) | nearest neighbor | L_HIP-body | 0.046 |
| Reappraisal (negative) | nearest neighbor | LH_SalVentAttn_FrOperIns_1 | 0.044 |
| Reappraisal (negative) | nearest neighbor | RH_Cont_PFCv_1 | 0.044 |
| Reappraisal (negative) | nearest neighbor | L_THA-intralaminar | 0.043 |
| Reappraisal (negative) | nearest neighbor | RH_Limbic_TempPole_1 | 0.043 |
| Reappraisal (negative) | nearest neighbor | LH_Default_PHC_1 | 0.043 |
| Reappraisal (negative) | nearest neighbor | LH_SomMot_4 | 0.043 |
| Reappraisal (negative) | nearest neighbor | RH_Default_Temp_1 | 0.042 |
| Reappraisal (negative) | nearest neighbor | R_PUT-anterior | 0.041 |
| Reappraisal (negative) | nearest neighbor | R_GP-internal | 0.041 |
| Reappraisal (negative) | nearest neighbor | L_AMY | 0.040 |
| Reappraisal (negative) | nearest neighbor | LH_Limbic_OFC_1 | 0.039 |
| Reappraisal (negative) | nearest neighbor | LH_Default_PFC_2 | 0.039 |
| Reappraisal (negative) | nearest neighbor | LH_SomMot_11 | 0.038 |
| Reappraisal (negative) | nearest neighbor | LH_Default_PFC_4 | 0.038 |
| Reappraisal (negative) | nearest neighbor | RH_Cont_PFCmp_1 | 0.037 |
| Reappraisal (negative) | nearest neighbor | RH_SomMot_8 | 0.037 |
| Reappraisal (negative) | nearest neighbor | RH_Default_Temp_2 | 0.037 |
| Reappraisal (negative) | nearest neighbor | RH_Default_pCunPCC_1 | 0.037 |

Note: [a] The cortical labels corresponding to cortical parcellations are from 200-region version of Schaefer parcellation atlas (Schaefer et al., 2018), meanwhile the subcortical labels corresponding to subcortical parcellations are from 32-region of Melbourne Subcortex Atlas (Tian et al., 2020).
